# Tmem138, a photoreceptor connecting cilium (CC) protein, is required for rhodopsin transport across the cilium and outer segment (OS) biogenesis

**DOI:** 10.1101/2021.05.22.445289

**Authors:** Dianlei Guo, Jiali Ru, Lijing Xie, Mingjuan Wu, Yingchun Su, Shiyong Zhu, Shujuan Xu, Yanhong Wei, Xialin Liu, Yizhi Liu, Chunqiao Liu

## Abstract

Photoreceptor connecting cilium (CC) is structurally analogous to the transition zone (TZ) of primary cilia and gates the molecular trafficking between the inner and the outer segment (OS). Retinal dystrophies with underlying CC defects are manifested in a broad array of syndromic conditions known as ciliopathies as well as non-syndromic retinal degenerations. Despite extensive studies, protein trafficking across the photoreceptor CC is largely unknown. Here we genetically inactivated mouse *Tmem138*, a gene encoding a ciliary membrane protein localized to the ciliary TZ and linked to Joubert syndrome (JBTS). Germline deletion of *Tmem138* abolished OS morphogenesis followed by rapid photoreceptor degeneration. Tmem138 was found localized to the photoreceptor CC and, accordingly, the molecular compartments of the CC and axoneme of the mutant photoreceptors were altered despite ciliogenesis proceeding normally at the early stage of photoreceptor development. To gain further insights into Tmem138 function in OS biogenesis, we focused on trafficking of rhodopsin, the most abundant protein of the OS. Mislocalization of rhodopsin was readily observed as early as P5 in the mutant photoreceptors prior to growth of the OS. Ablation of *Tmem138* in mature rods recapitulated the molecular changes in the germline mutants, causing well-formed outer segment discs to disintegrate accompanied by mislocalization of rhodopsin in the cell body. Furthermore, Tmem138 interacted with rhodopsin, and two additional CC compartment proteins Ahi1 and Tmem231, which were both altered in the mutant photoreceptors. Taken together, these results suggest that Tmem138 has a distinct role in gating the transport of rhodopsin and likely other OS bound proteins through formation of CC transport complex(es).

## Introduction

Ciliopathies are pleiotropic genetic diseases affecting multiple organs, and can be clinically classified as distinct entities including Bardet–Biedl syndrome (BBS), Meckel-Gruber syndrome (MKS), Senior Loken syndrome (SLS), and Joubert syndrome (JBTS) among others. The photoreceptors of the retina are affected in nearly all ciliopathies as their light sensing outer segments (OS) are modified cilia and thus highly vulnerable. Approximately 10% of the OS is renewed each day, requiring active transport of a large amount of newly synthesized proteins and lipids across the cilium in support of OS renewal. Rhodopsin is the most abundant OS disc protein, mutations of which alone account for 10-20 % retinitis pigmentosa (RP). Vectorial transport of rhodopsin from the biosynthetic inner segment to the outer segment is crucial for both photoreceptor homeostasis and functions, and ectopic rhodopsin localization is a major cause of photoreceptor death (Grimm et al., 2000; Li et al., 1996; Louie et al., 2010; Sung et al., 1994). Despite extensive studies, rhodopsin transport has only been partially understood. Trafficking of rhodopsin-carrying vesicles from ER/Gogi to the base of the connecting cilia is coordinated by GTPases, their effectors and the intraflagellar transport (IFT) complexes (Keady et al., 2011; Pearring et al., 2013; Wang and Deretic, 2014). Recent evidence from live-imaging in both cultured cells and retinal explants suggested that IFT88 and Kif3a were involved in ciliary transport of rhodopsin (Trivedi et al., 2012), presumably along the axonemal microtubules. Although many mouse models with mutations in IFT subunits or associated motors exhibit mislocalization of rhodopsin (Marszalek et al., 2000; Pazour et al., 2002), the causality between these transporting components and rhodopsin trafficking defect is inconclusive, often due to ciliary and/or OS structural defects of the mutant photoreceptors. Furthermore, aside from the proposed general protein transport machinery, whether ciliary membrane components are also involved in rhodopsin trafficking has not been reported.

The photoreceptor outer segment is a modified sensory cilium that shares common structural features with that of other cells (Wang and Deretic, 2014). At the base, it grows from the basal body (BB) derived from a mother centriole, and further extends ciliary axonemes (Ax) distally, which are microtubule arrays supporting OS membrane discs and providing routes for IFT. The most proximal zone of the Ax is known as the connecting cilium (CC), structurally equivalent to the ciliary transition zone (TZ) of non-photoreceptor cells. The connecting cilia and associated membranous structures function as the ciliary gate/diffusion barrier and plays a crucial role in selecting targeting proteins allowed to enter the outer segments (Leroux, 2007; Nachury et al., 2010; Wang and Deretic, 2014). Genetic studies in *C. elegans* and in vertebrate models have defined two TZ modules, the NPHP and MKS modules. The NPHP module includes NPHP-1, NPHP-4 and NPHP-5 (IQCB1) (Barker et al., 2014; Winkelbauer et al., 2005), and MKS module contains MKS-1, MKSR-1 (B9d1), MKSR-2 (B9d2), MKS-2 (TMEM216), MKS-3 (TMEM67/Meckelin), MKS-6 (CC2D2A), TMEM231, JBTS-14 (TMEM237), TMEM107, TCTN-1 and AHI1/Juberin (Barker et al., 2014; Li et al., 2016; Williams et al., 2011). Both modules are assembled on MKS5 (CEP290)/RPGRIP1L scaffold (Li et al., 2016). Interestingly, two additional components TMEM138 and CDKL-1, genetically dependent on MKS5/CEP290 but no other MKS or NPHP proteins, are considered as another potential distinct module (Jensen et al., 2015; Li et al., 2016).

In the mammalian system, much has been learned about Rpgr/Cep290-centered interactome including these of NPHP and BBS proteins (Baehr et al., 2019; Rachel et al., 2012; Rachel et al., 2015); On the other hand, the MKS TZ module is relatively understudied in higher organisms, especially for a group of TZ-localized putative transmembrane proteins (TMEM17, 67, 107, 138, 216, 218, 231, 237) (Collin et al., 2012; Garcia-Gonzalo et al., 2011; Huang et al., 2011; Jensen et al., 2015; Lambacher et al., 2016; Li et al., 2016; Roberson et al., 2015), the localization and ciliary function of which have been scarcely explored in either humans or mice. Thus, whether the data obtained from the lower organisms or in vitro cultured cells resembles that of mammals has yet to be verified.

In our attempts to investigate the potential roles of planar cell polarity (PCP) signaling in photoreceptor morphogenesis, we identified several genes coding for transmembrane proteins with altered expression in the *Prickle 1* (a core PCP component) mutant retinas, including *Tmem138* and *Tmem216* known to underlie JBTS (Lee et al., 2012; Suzuki et al., 2016; Tuz et al., 2013). *Tmem138* and *Tmem216* are arranged in a head-to-tail configuration on human chromosome 11 and mouse chromosome 19, sharing intergenic regulatory elements (Lee et al., 2012). Tmem138 is localized at the ciliary base and axoneme, whilst Tmem216 is primarily in the basal bodies in IMCD3 cells (Lee et al., 2012). Tmem138 was suggested to coordinate interdependent vesicular transport to the primary cilia in vitro, and its human mutations reduce cilia length of the patient fibroblasts (Lee et al., 2012). To understand their functions in photoreceptor OS morphogenesis, we genetically deleted *Tmem138* in mice in the present study and report the complete failure of OS development in the mutants. Tmem138 also interacts with rhodopsin and two CC-localized proteins, Ahi1 and Tmem231, which altogether might be a part of OS protein transporting machinery across the cilium.

## Materials and Methods

### Animal husbandry

All procedures involving the use of mice were approved by Zhongshan Ophthalmic Center Animal Care and Use Committee (ACUC). Transgenic mice were generated using Cyagen service (www.cyagen.com).

### Generation of Tmem138 mutant alleles

The ‘knockout-first’ strategy was used to create the *Tmem138* gene-trap allele (*Tmem138^a/a^*) by homologous recombination in ES cells (Fig. 1A) (Liu et al., 2013; Testa et al., 2004). Briefly, a gene-trap construct was engineered with a Frt site and *En2* splicing acceptor (SA) preceding an *eGfp* reporter, followed by a polyadenylation signal, *loxP*, *PGK-neo*, *Frt*, and *loxP*. This gene-trap cassette was placed between *Tmem138* exon 1 and 2. A third *loxP* was placed right after exon 3, which will be used for the generation of a conditional knockout allele. Southern analysis was performed as described previously (Liu et al., 2013) to identify the F1 *Tmem138^a/a^* allele. A 5’-probe outside of the left arm was used to identify 8.3kb- and 12.4kb-ApaLI fragments from wild type and *Tmem138^a/a^* alleles, respectively (Fig.1B).

**Figure 1.**
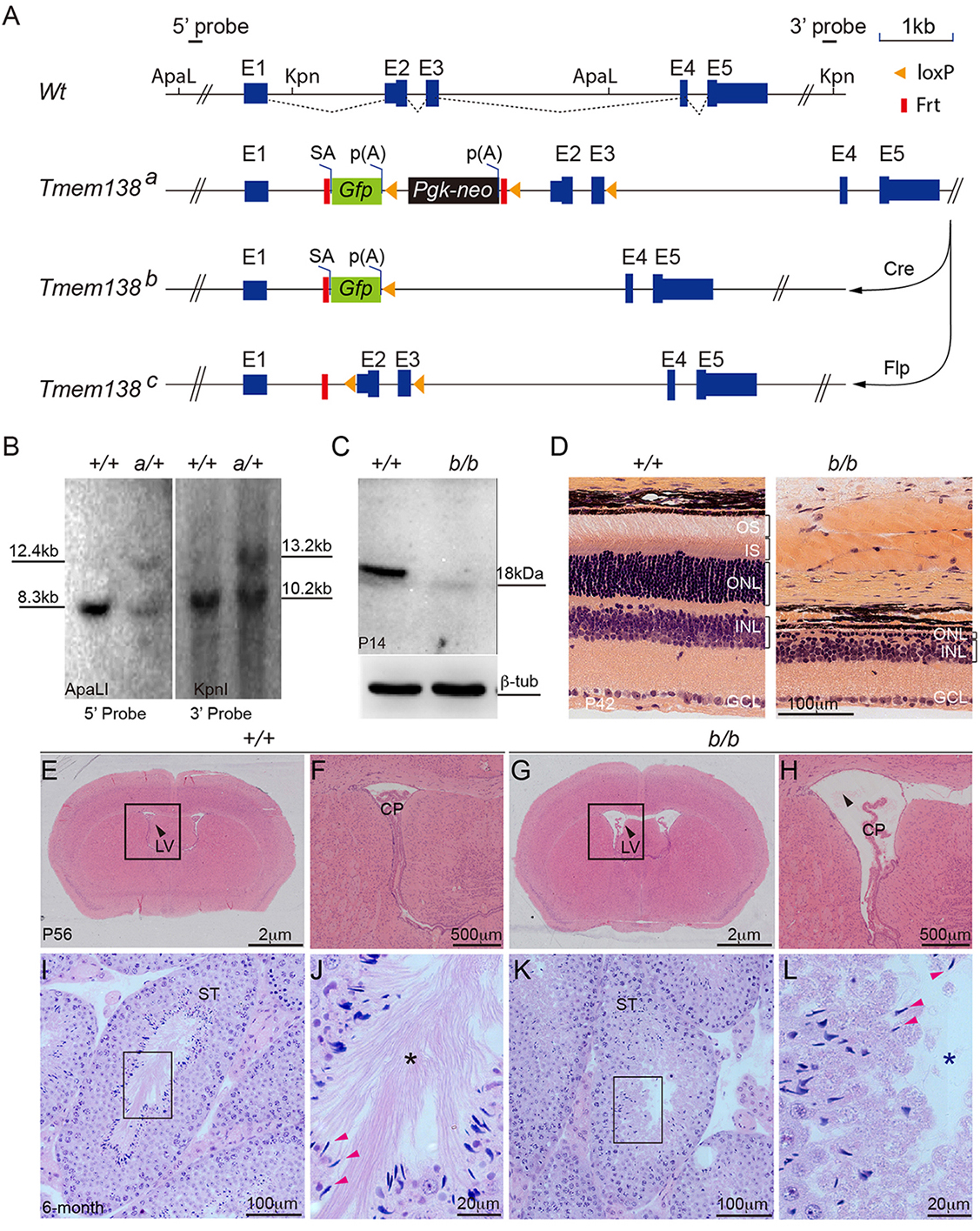
Targeted inactivation of *Tmem138* leads to enlarged brain ventricles, azoospermia, and retinal degeneration. (A), *Tmem138* targeting strategy. The *Tmem138* gene has five exons with protein coding starting at Exon 2 (E2). The restriction enzyme sites ApaLI and Kpn1 were used for Southern blotting analysis in (B) with 5’ and 3’ probes, respectively. SA, splicing acceptor; p(A), polyadenylation signal; *Gfp*, green fluorescent protein; *Pgk-neo,* phosphoglycerate kinase promoter (*Pgk*)-driven neomycin gene (neo). (B), Southern analysis of *Tmem138* a gene trap allele (*Tmem138 ^a^*). The expected fragment sizes on Southern blot are 5’ Probe (ApaLI): 8.30 kb (*Wt: +/+*) & 12.41 kb (*Tmem138 ^a^ :a/+*); and 3’ Probe (KpnI): 10.26 kb (*+/+*) & 13.28 kb (*a/+*), respectively. (C), Western blotting of P14 retinal extracts using an anti-Tmem138 antibody. (D), Homozygous *Tmem138^b/b^* (*b/b*) null mutant manifested severe retinal degeneration at postnatal day 42 (P42). OS, outer segment; ONL, outer nuclear layer; INL, inner nuclear layer; GCL, ganglion cell layer. (E), A coronal brain section from wild-type mouse stained with Hematoxylin and Eosin (H&E) at P56. Arrow points to the lateral ventricle (LV). Boxed area is magnified in (F). CP; Choroid plexus. (G), Enlarged brain ventricles of the ‘*b/b*’ mice. Boxed area is magnified in (H). (I, K), Testis sections from 6-month-old wild type and null mutant mice, respectively. Boxed areas show the lumen of seminiferous tubes (ST) and are magnified in (J, L). Red arrows point to sperm heads; asterisks indicate sperm tails, which are well developed in the ‘*+/+*’ but absent from the *‘b/b’* mice.

For the generation of *Tmem138^b/b^* straight knockout allele, *Sox2-Cre* transgenic females (equivalent to the germline *Cre* transgenic lines, JAX laboratory) were used to cross with the *Tmem138^a/a^* to delete exon 2 and 3, and leaving the *eGfp* reporter driven by the endogenous *Tmem138* promoter. For the generation of the *Tmem138^c/c^* conditional allele, the *CAG-Flp* transgenic line was used to delete the gene-trap cassette, leaving two loxP sites flanking the exon 2 and 3 (Fig. 1A). Further crossing *Rho-Cre* transgenic line with the *Tmem138^c/c^* allele generated rod-specific *Tmem138* mutants.

### Genotyping and detection of *Tmem138* allelic transcripts and proteins

All primers used in this study were listed in Suppl. Table 1. Primers for genotyping and detection of *Tmem138* allelic transcripts were illustrated in Suppl. Fig. 1 & 7. Briefly, primer pairs of p1-p2 and p3-p4 were used for genotyping of wild type and *Tmem138 ^b/+^* straight knockout allele (Tm138 b), respectively. Primer pair p13-p14 was for genotyping of both wild type and *Tmem138 ^c/+^* conditional knockout allele; and primer pair p15-p16 was for genotyping *Rho-Cre* transgenic allele. PCR programs for p1-p2, p3-p4, p13-p14 and p15-p16 are all the same as follows 94℃ 5min; 94℃ 10s, 61℃ 30s, 72℃ 30s, 35 cycle; 72℃ 10min.

For RT-PCR, retinas from wild type and *Tmem138* mutant mice at postnatal 12 days were dissected in PBS on ice. Fresh retinas were subjected to RNA extraction using RNAsimple Total RNA Kit (DP419, TIANGEN, China) per the manufacturer’s instructions. Total RNAs were reverse transcribed into cDNAs (A5001, Promega, China), and PCR reactions were performed to identify transcripts from different *Tmem138* alleles. P5-p6, p7-p8, p9-p10, and p11-p12 were used for genotyping exon (E) 1, *Gfp*, E2, and E5, respectively. PCR programs for p5-p6, p7-p8, p9-p10 and p11-p12 are all the same as follows 94℃ 5min; 94℃ 10s, 60℃ 30s, 72℃ 30s, 35 cycles; 72℃ 10min.

Southern blotting was performed as described previously (Liu et al., 2013). Briefly, α^32^P-labeled probes outside the left (5’-probe) and the right (3’-probe) homologous arms were hybridized to nylon membranes carrying ApaLI- and KpnI-digested wild type and *Tmem138* ^a/a^ genomic DNA, respectively. The 5’-probe detected 8.3kb wild type and 12.4kb mutant fragments; and the 3’-probe detected 10.2kb wild type and 13.2kb mutant fragments. The Southern probe primer sequences are provided in Suppl. Table 1.

For Western blot analysis, protein samples were blotted onto polyvinylidene difluoride (PVDF) membranes by a wet transblot system (Mini-Protein Tetra; Biorad) using the standard protocol recommended by the manufacturer. Antibody against Tmem138 (HPA042373; Sigma, Suppl. Table 2) was used to probe the blot using a standard protocol. Chemiluminescent images were taken by FluorChem R (Proteinsimple).

### Histology, immunohistochemistry, and TUNEL labeling

For plastic sectioning, tissues (eyeball, brain, cerebellum, kidney, testis) were dissected and fixed with Carnoy’s fixative (60% alcohol, 30% chloroform,10% Acetic acid) overnight at room temperature, dehydrated with a graded series of ethanol, and embedded using Technovit kit according to the manufacturer’s instruction (14653, Technovit, Germany). 3 µm plastic sections were cut with a microtome (RM 223, Leica, Germany) and stained with hematoxylin and eosin according to the manufacturer’s instruction (UI0402, UBIO, China).

Immunohistochemistry and TUNEL (Roche, Cat. 11684795910) detection of apoptosis were performed as described previously by Guo et. Al (Guo et al., 2018). Antibodies were listed in Suppl. Table 2 with vendor catalog numbers and experimental conditions.

### Optical coherence tomography (OCT) and Electroretinography (ERG)

For full-field ERG testing, mice were dark-adapted overnight and anesthetized by an intraperitoneal injection of 4-5 µl/g 10% chloral hydrate. Pupils were dilated using Compound Tropicamide Eye Drops (SINQI, China). ERG responses were recorded using CELERIS system (D430, Diagnosys LLC, USA) with a stimulating electrode in contact with the cornea through Hypromellose eye drops (ZOC, China). Scotopic and photopic ERGs were recorded using strobe flash stimuli from 5-averaged responses of 0.003 ∼10cd.s/m^2^ and 0.3∼30 cd-s/m^2^, respectively. Light adaptation with background intensity at 30 cd/m^2^ was performed for 5 minutes between scotopic and photopic recordings. The values of a- and b-waves were exported to GraphPad Prism 5 for statistical analysis. Student *t-tests* were performed to determine statistical significance. A total of 6 animals were used for each genotype.

For OCT, anesthetization and pupil dilation were performed as described for ERG recording. OCT images were acquired using a Spectralis HRA + OCT system (Heidelberg Engineering, Heidelberg, Germany). Mice were placed on a stage for alignment and corneas were kept moist with drops of 0.9% sterile saline. Images were taken centered on the optic nerve head, and horizontal or radial scans were obtained from each eye. The thickness of the outer retina (RPE + OS + ONL +OPL) and the inner retina (INL + IPL +GCL) were analyzed using Heidelberg Spectralis system software, and values of a- and b-waves were exported to GraphPad Prism 5 for statistical analysis. Stuent *t-test* was performed to obtain the statistic powers. A total of 6∼10 animals were used for each genotype.

### Transmission electron microscopy (TEM), scanning electron microscopy (SEM), and immuno-electron microscopy (Immuno-EM)

For both TEM and SEM, P0∼P14 mice were euthanized by decapitation or cervical dislocation as described previously (Guo et al., 2019). Eyes were enucleated and immersed in 2%PFA for 10 min, and the retinas were dissected out, cut into small pieces (about 2mm x 1mm) and fixed further in 2% PFA/2.5%/glutaraldehyde in PBS at 4℃ overnight. Adult mice were deeply anesthetized with 10% chloral hydrate (5ul/g) and cardiac perfusion was performed first with 50 ml 0.9% saline followed by 50 ml of fixative (2% PFA, 2.5% glutaraldehyde in PBS) for about 10 min. The enucleated eyes were further fixed in the same fixative for 24 h and cut on a vibratome (VT1000S, Leica, Germany) at 200μm thickness.

For TEM, retina pieces or sections were then rinsed with 50 mM cacodylate buffer (PH 7.4) and incubated in 1% reduced OsO4 in cacodylate buffer on ice for 90 min. Sections were dehydrated through a gradient series of ethanol, infiltrated with propylene oxide, and embedded in EMBED812 resins (14120, EMS, USA) at 60℃ for 36-48 hours. Ultrathin sections at 60-80 nm were cut on an ultramicrotome (Leica EM UC7, Leica, Germany) and collected onto copper grids (FCF200-Cu-50, Electron Microscopy Sciences, USA). Sections were poststained with 1% uranyl acetate for 30min and Sato’s lead for 3min. Electron microscopy images were acquired on HT-7700 (Hitachi, Japan).

For SEM, the cornea, sclera, and choroid were quickly removed from the freshly dissected eyeballs. Retinas with lens and RPE were then fixed in the same fixative as for TEM. Vibratome sections were prepared at 200 μm thickness after removal of lens, dehydrated with ethanol and acetone. The samples were then immersed in isoamyl acetate, dried, sputter-coated, and imaged using scanning electron microscopy (S-3400N, Hitachi, Japan).

For immuno-EM, P5, P8, and P14 retinas were cut into small pieces about 2mm x 1mm after 2%PFA fixation. Retina pieces were fixed in 2% PFA/0.1% glutaraldehyde in PBS at 4℃ overnight. The retina pieces were rinsed with PBS, dehydrated with ethanol, and embedded in capsules filled with LR white resins (14381-UC, EMS, USA). The resins were let to polymerize at 60℃ for 36-48 hours in an oven. 60-80nm ultrathin sections were cut and collected onto nickel grids (FCF200-Ni-50, EMS, USA). Sections were treated with 50 mM Tris glycine buffer (pH7.4) for 10 min, rinsed with PBS, and blocked with 10% BSA in PBS at room temperature (RT) for 30 min. After incubating overnight at 4℃ with 1D4 antibody (anti-rhodopsin C terminus), grids were washed with 1% BSA in PBS for 5 times, 5min each, and probed with IgG-conjugated 10nm colloidal gold particles (25825, EMS, USA) for 2h at RT. Grids were rinsed and counterstained with 1% uranyl acetate for 5-10min and Sato’s lead for 30s-1min.

### Detection of Protein interaction

Plasmid constructs: pLVX-*3F-TST-Tmem138*: 3xFlag and Twin-Strep-tagged (TST) tags at the N-terminus; pRK5 *HA-rhodopsin*: HA tag at the N-terminus; pRK5-*Ahi1-HA/Flag* and pRK5-*Tmem231-HA/Flag*: HA or Flag tag at the C-terminus; pRK5-*Tmem67-Flag,* pRK5-*Arl13b-Flag,* and pRK5-*Rab8a-Flag*: Flag tag at the C-terminus. Construct diagrams are listed in Suppl. Fig. 11A.

HEK293T cells were co-transfected with equimolar amounts of the tagged plasmids (total 10ug/100mm^2^ dish, ∼7×10^6^ cells) and were harvested after 48 h in culture. Briefly, cells were incubated on ice for 15min in 1ml of TBS lysis buffer containing 30mM Tris-HCl (PH 7.4), 150mM NaCl, 0.5%NP-40 (I8896, Sigma, USA) and a protease inhibitor cocktail (P8340, Sigma, USA). The lysed cells were centrifuged for 15 min (10,000xg at 4°C). For 3F-TST-tagged proteins, the supernatant was incubated with 100μl Strep-TactinXT resins (2-4010-010, IBA, Germany). For HA- or Flag-tagged proteins, the supernatant was incubated with 100μl Anti-HA Affinity gel (D111140, Sangon Biotech, China) or Anti-Flag M2 Affinity Gel (A2220, Sigma, USA). Incubation lasted overnight at 4℃ with rotation. The mixtures were then loaded into empty chromatography columns, and resins or agarose beads were packed by gravitation. Columns were washed 3 times withTBS buffer to clear off residual unbound proteins and eluted with 3 times 100μl biotin for TST column, or HA, or Flag peptides for Sepharose columns. The eluant from each fraction was examined by Western blotting using anti-Flag or anti-HA antibody. The eluant having the maximal amount of bait proteins was used for detection of protein interaction. Schematic illustration of affinity purification is shown in in Suppl. Fig. 11B.

For retinal extracts immune-pull-down experiments,14 retinas from 3-week-old mice were dissected and snap frozen in liquid nitrogen. Retinas were then transferred into TBS lysis buffer (30mM Tris-HCl (PH 7.4), 150mM NaCl, 0.5%NP-40) supplemented with protease inhibitor cocktail (P8340, sigma, USA) and gently vortexed for 10 seconds. The OS and CC components were separated into supernatant from the retinal tissue chunks by spinning down at 2000G for 10S. Let the supernatant stayed in the lysis buffer for another 15 min on ice, and centrifuged at 15000 x g at 4°C to remove the tissue debris.

For protein pull-down assays, 100 μl/sample Pierce protein A/G magnetic beads (88803, Thermo, USA) were blocked with 10% BSA at 4℃ overnight on a rotary plate. The above-prepared OS and CC protein extracts were incubated with Tmem138 antibody at 4°C overnight with rotating. The sample/antibody mixtures were transferred to a 1.5mL microcentrifuge tube containing the pre-blocked magnetic beads and incubated at room temperature for 1 hour with rotating. The magnetic beads were then collected with a magnetic stand, and proteins were eluted with 100μl of glycine (50 mM, pH2.0) and analyzed by SDS-PAGE electrophoresis and Western blotting.

### Quantification and statistics

For quantification of the photoreceptor ciliary structures, TEM images from P3 and P5 retinas were acquired as previously described. Cilia between RPE and the ONL at each stage of photoreceptor ciliogenesis were counted from 3 retinal sections/retina, each longitudinally spanning over a 100μm length in mid-peripheral retinal areas. The numbers of cilia from 3 sections (3×100 μm=300 μm total span) for each stage were used for plotting the graphs. One retina for each genotype was counted.

For measuring ciliary density, Ac-tub stained flat-mount retina was divided into three areas: central, mid-peripheral, and peripheral. Images were taken with a 40x/1.40 oil objective at 1024 x 1024-pixel resolution. The cilia numbers were determined by counting the number of cilia in the above three areas using image J and normalized to the counting area (per 100 μm^2^).

To measure lengths of ciliary domains stained with immunofluorescent markers, images were taken with an 100x/1.40 oil objective and 3.0x optical zoom factor at 1024 x 1024 resolution, using Zeiss LSM880 equipped with Airyscan. A total of 100∼150 cilia of each retina section/animal (total 3 animals) were used. The lengths of stained ciliary domains were determined by Image J software (National Institutes of Health).

Tmem231 fluorescence intensities in the CC were determined as fluorescent signals/CC area with ImageJ. A total of 100∼150 cilia of each retina section/animal (total 3 animals) were used for the quantification.

All statistical analyses were performed with the GraphPad Prism 7.0 software and presented as mean ± SD. Statistical powers were determined by the Student’s t-test. ‘*’, p < 0.05; ‘**’, p< 0.01.

### Tmem138 tertiary structure prediction

I-TASSER (Iterative Threading ASSEmbly Refinement) on-line service from Zhang Lab were used for Tm138 protein 3D structure predication. (https://zhanglab.ccmb.med.umich.edu/I-TASSER/), It first identifies structural templates from the protein database (PDB) by multiple threading approach LOMETS, with full-length atomic models constructed by iterative template-based fragment assembly simulations. Functional insights of the target are then derived by re-threading the 3D models through protein function database BioLiP.

## Results

### Targeted inactivation of *Tmem138* leads to enlarged brain ventricles, azoospermia, and retinal degeneration

An early investigation on planar cell polarity (PCP) in mouse retina identified several genes coding for transmembrane proteins and regulated by *Prickle 1,* a core PCP component (data not shown). One of them is *Tmem138*, mutation of which causes Joubert syndrome spectrum disorder affecting multiple organs including the brain, testis, kidney, and retina (Lee et al., 2012; Suzuki et al., 2016). To understand the mechanisms of retinal dystrophy in JBTS caused by *TMEM138* mutations, we genetically inactivated the *Tmem138* gene using a Knockout-first strategy in mice (Fig. 1). A *Gfp* reporter with a splicing acceptor (SA) and a polyadenylation (p(A)) signal followed by a selection cassette *Pgk-neo* was inserted between exon 1 (E1) and 2 (E2) to create a gene-trap allele (*Tmem138 ^a^*). *loxPs* and *Frts* were arranged such that excision with Cre or Flp recombinases would generate a straight or a conditional knockout allele, respectively (Fig.1A). The gene-trap allele was confirmed by Southern blotting analysis of mouse tail DNA (Fig. 1B). A germline Cre (*Sox2-Cre*, see Materials and Methods) was crossed onto *Tmem138^a^* to delete coding exons 2 and 3 and the *Pgk-neo* cassette to create a null allele (*Tmem138^b^*) (Suppl. Fig. 1A, B). Targeted RT-PCR was conducted to probe exonal transcripts from each allele (Suppl. Fig. 1A, C, D). Notably, *Tmem138^a^* allele appeared to transcribe all exons indicated by existence of the last exon E5 transcripts, whereas *Tmem138^b^* allele only expressed E1 and knock-in *Gfp* reporter (Suppl. Fig. 1C, D) indicating its nullity. Western blotting further confirmed the loss of Tmem138 protein in *Tmem138^b/b^* mice (Fig. 1 C). *Tmem138 ^b/b^* mutants had a normal appearance and similar growth curves with that of wild type mice (Suppl Fig. 1E), and numbers of mice born for each genotype from *Tmem138 ^b/+^* breeding approximately adhered to the Mendelian ratio (Suppl Fig. 1F).

We next performed histological examinations of tissue sections using hematoxylin and eosin (H&E) staining. Rapid photoreceptor loss was observed in the mutant retina with only one row of nuclei remaining at P42 (Fig. 1D). Mutant mice also exhibited hydrocephalus as evidenced by enlarged brain ventricles (Fig. 1E-H) and defective spermatozoa (Fig. 1I-L). The homozygous gene-trap mice (*Tmem138 ^a/a^*) exhibited the same phenotypes as that of the *Tmem138 ^b/b^* null mice (data not shown). Western blotting also verified the absence of Tmem138 protein in the *Tmem138 ^a/a^* mutants (data not shown), probably due to a complete blocking of the *Tmem138* mRNA translation from the stop codon of *Gfp* or *neo*. All noted phenotypes were fully penetrant in the examined animals. Thus, the *Tmem138* mutants manifested a subset of the JBTS phenotypes.

### Early-onset rod loss in *Tmem138* mutants

Severe retinal degeneration of the mutant mice within 6 weeks of birth indicates an early-onset photoreceptor loss. To further clarify the temporal windows of photoreceptor loss upon *Tmem138* disruption, we performed optical coherence tomography (OCT) in live mice to measure the retinal thickness of *Tmem138* mutant mice and wild type littermates. Significant thinning of the mutant outer retina (OS + ONL+ OPL) (∼20μm) was readily detectable as early as P14 (Fig. 2A, Suppl Fig. 2A, B). Rapid loss of the ONL occurred in the ensuing weeks, with the outer retinal thickness reduced to about 50% (~ 50 μm) by P21 (Fig. 2B, Suppl Fig. 2C, D). Continuous but slower thinning of ONL was detected till complete loss of ONL at P42 (Fig. 2C, Suppl. Fig. 2E, F). A reduction of the inner nuclear layer (INL) was also observed at the examined ages but was much less severe compared to that of the outer retina (Fig. 1 D, Suppl. Fig. 2B, D, F). Consistent with the thinning ONL, electroretinography (ERG) measuring photoreceptor light responses revealed that the mutant rod response was essentially abolished at P21, whereas the cone response was diminished but still measurable under higher light intensities (Fig. 2D, Suppl. Fig. 2G). By P42, both rod and cone responses were undetectable (Fig. 2E, Suppl. Fig. 2H).

**Figure 2.**
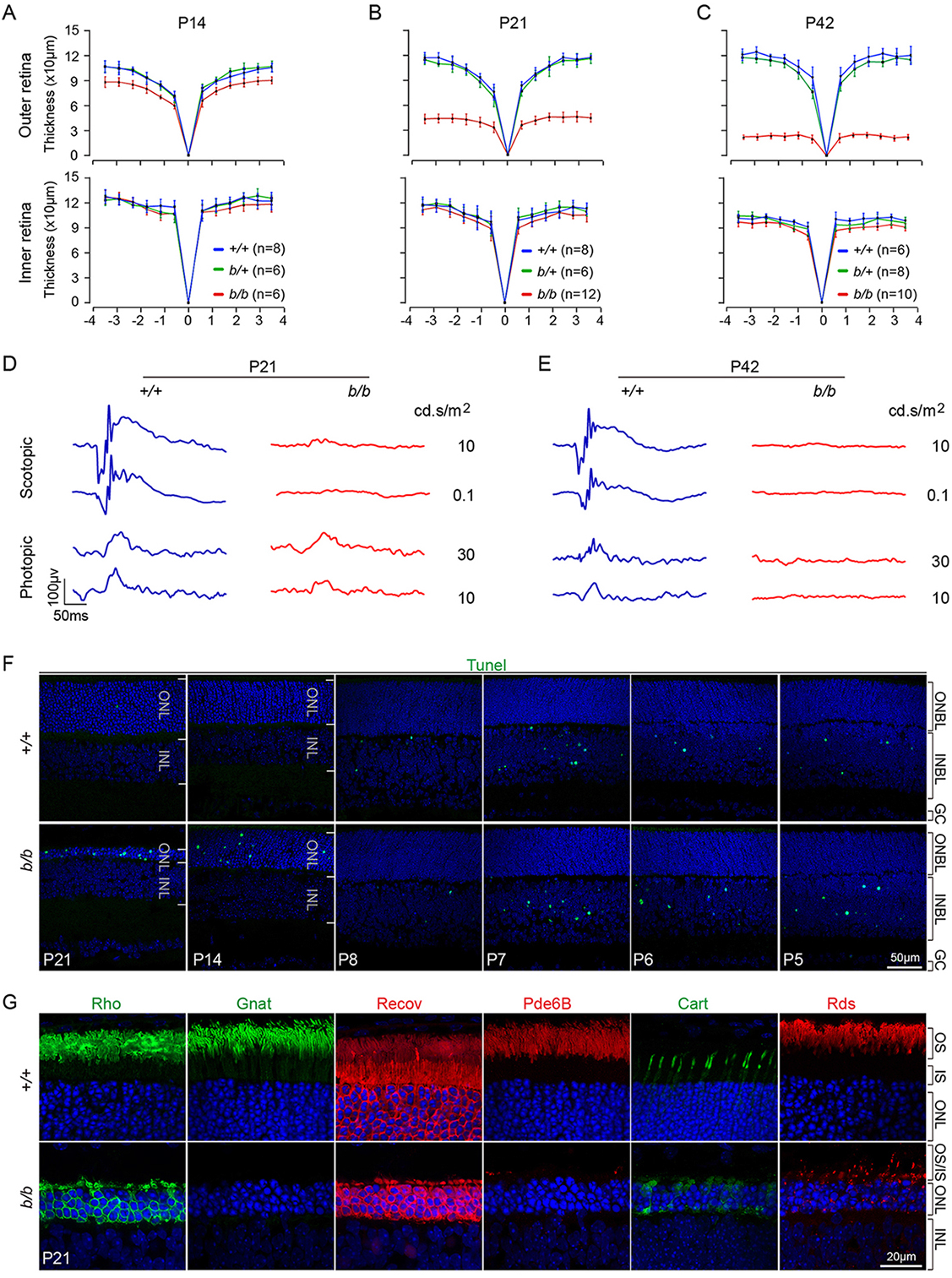
Structural and functional retinal defects of *Tmem138^b/b^* mice. (A-C), Optical coherence tomography (OCT) imaging analysis of retinal thickness at different ages. (D, E), Retinal electroretinography (ERG) at P21 and P42. (F), Detection of cell death by TUNEL labeling at indicated ages. Note apoptosis at early ages from P5 to P8 in the INL are normal during retinal maturation. (G), Immunostaining with antibodies against rhodopsin (Rho), alpha transducin (Gnat), Recoverin (Recov), phosphodiesterase 6B (Pde6B), cone arrestin (Cart), and Peripherin/RDS (Rds).

To further corroborate the findings of OCT and ERG, we examined cell death at different ages of photoreceptor development. Extensive cell death was detected at both P14 and P21 by TUNEL labeling (Fig. 2F, Suppl. Fig. 2I). A time series of investigations revealed that cell death started between P8 and P14 (Fig. 2F). Additionally, staining of Caspase-3, TUNEL & phospho-Histone 2A isoform X, p-H2A.X (for DNA damages), and LCA3A/B (for autophagy) was all elevated as early as P14 (Suppl. Fig. 2J). Muller glial response to retinal injury was conspicuous at P21 as indicated by upregulation of GFAP (Suppl. Fig. 2K).

We next examined several key phototransduction proteins by immunostaining in mutant and control retinas at P21 (Fig. 2G). Rhodopsin (Rho), transducin alpha (Gnat1), recoverin (Recov), and phosphodiesterase 6B (Pde6B) were all barely detected from the mutant rod outer segment (OS), with a considerable fraction of rhodopsin mislocalized in the cell bodies (Fig. 2G). In contrast, the numbers of cones labeled with cone arrestin (Cart) were comparable between the mutant and control retinas, although mislocalization of Cart was seen throughout cell bodies of mutant photoreceptors (Fig. 2G). Outer segment disc protein peripherin 2 (Rds) essentially disappeared from the mutant OS, with punctate staining remaining throughout the cell bodies (Fig. 2F). Thus, the data together demonstrated an early-onset photoreceptor degeneration upon disruption of *Tmem138*.

### Localization of Tmem138 to the connecting cilium

To gain molecular insights into Tmem138 function in photoreceptors, we examined its protein localization. Previous studies suggest that Tmem138 is a ciliary protein in mouse IMCD3 cells (Lee et al., 2012) and localized in TZ in *C. elegans* sensory neurons (Li et al., 2016). It remains unclear about its localization in photoreceptors. Using the same antibody validated by Western blotting in Figure 1C, we performed immunohistochemistry on retinal sections in combination with an antibody against acetylated α-tubulin (Ac-tub), which labels a broad ciliary microtubule domain including the CC. Under light fixation (Materials and Methods), Tmem138 fluorescent signals appeared to be in the proximal region of the wild type CC but not the mutant (Fig. 3A-C, arrows). To further define the subcellular domain of Tmem138 localization, we performed immunohistochemistry in the adult photoreceptors. Tmem138 was co-labeled, in order, with axonemal protein Rp1, which is localized distal to the CC (Fig. 3D), Ac-tub, spanning the CC (Fig. 3E), γ-tubulin for the basal body (BB), which is immediately proximal to the CC (Fig. 7F), and rootlet protein Rootletin, which is proximal to the BB (Fig. 3G). The location of Tmem138 staining relative to these ciliary proteins (Fig.3H) suggests it is a CC component. *Tmem138* encodes 162 amino acids having a theoretical 18.4 kDa molecular weight with 4 transmembrane domains predicted with I-TASSER program (https://zhanglab.ccmb.med.umich.edu/I-TASSER/)((Fig. 3I). Thus, the results suggest that Tmem138 is likely a CC membrane protein.

**Figure 3.**
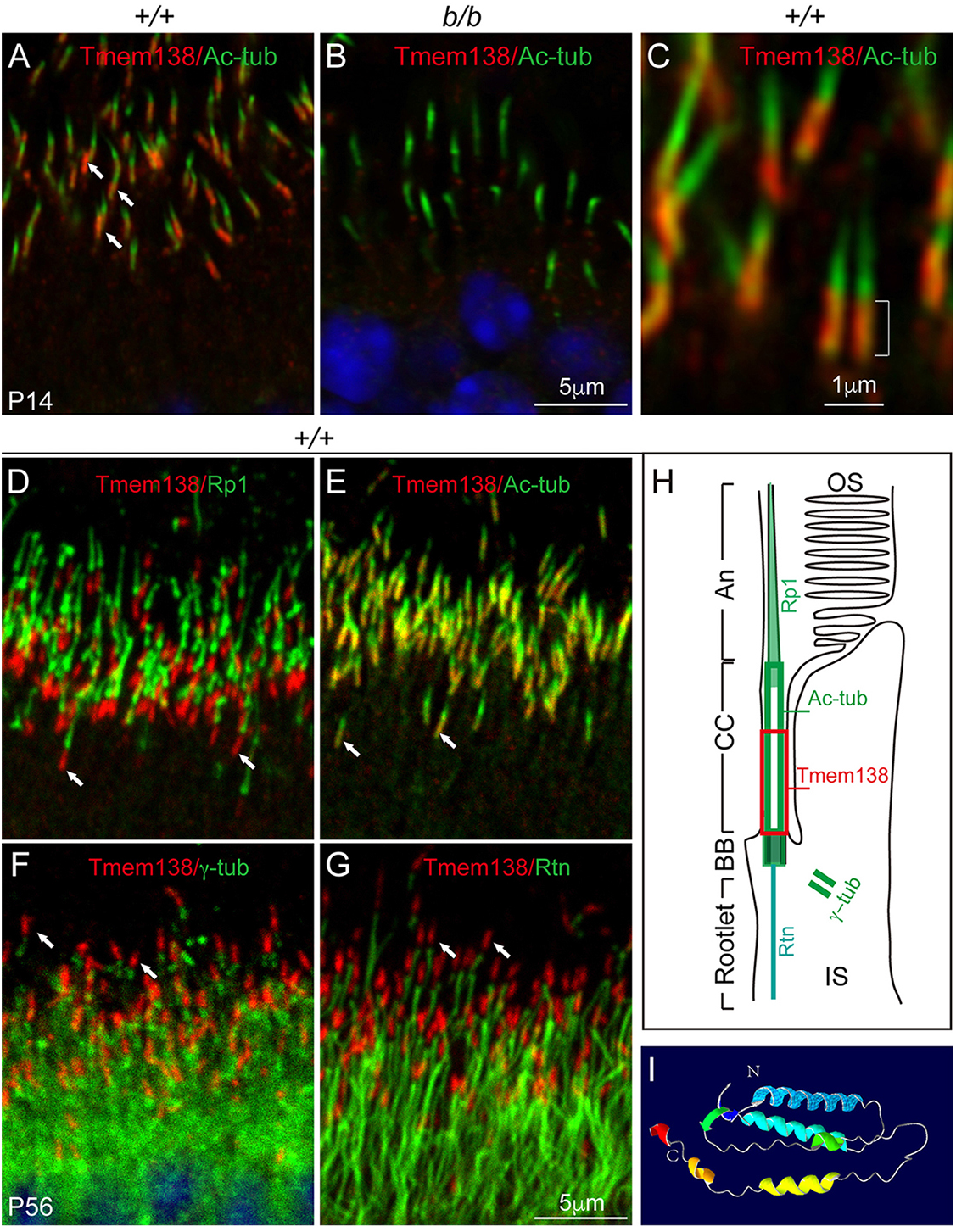
Tmem138 is localized to the connecting cilium. (A), Tmem138 was localized to the CC (arrows) at P14. (B), Tmem138 staining at the CC was abolished in the *‘b/b’* mutant. (C), Magnified cilia with Tmem138 and Ac-tubulin double staining at P14. A bracket indicates the proximal CC with Tmem138 staining (red). (D) to (G) are double staining of Tmem138 with Rp1 (D), Ac-tubulin (E), γ-tubulin (F), and Rootletin (G), respectively. Arrows point to Tmem138 staining (red). (H), A schematic drawing of a rod photoreceptor with the red boxed area indicating the Tmem138 localization compartment relative to that of other ciliary markers. (I), Bioinformatical prediction of Tmem138 with 4 transmembrane domains (I-TASSER, https://zhanglab.ccmb.med.umich.edu/).

### Altered molecular domains of CC and axonemal compartments of the mutant photoreceptors

The CC localization of Tmem138 prompted us to investigate whether photoreceptor ciliary compartments are altered upon disruption of *Tmem138*. Triple labeling with Ahi1, a CC compartmental protein, Ac-tub and Rp1, was performed on retinal sections of P5, P8, and P14. At P5, basal CC staining of Ahi1 (red) and axonemal staining of Rp1 (turquoise) was similar in wild type and the mutant retinas (Fig. 4A, B). A staining gap was present separating Ahi1 and Rp1 compartments at this age (Fig. 4A, B, brackets). Ahi1 staining domain extended distally becoming adjacent to that of Rp1 in about two thirds of cilia at P8 (Fig. 4C), whereas the gap between Ahi1 and Rp1 remained in most of the mutant cilia (Fig. 4D). Axonemal Rp1 extension was thwarted after P8 (Fig. 4. C-F), with further shortening of the Ahi1 domain in P14 photoreceptors (Fig. 4E, F). Quantification of the Ahi1 domain lengths and the staining gaps between Ahi1 and Rp1 demonstrated significant differences between wild types and the mutants (p<0.01) at P8 and P14, whilst the average length of Ahi1 domain plus the gap between Rp1 and Ahi1 was statistically unaltered (Fig. 4G). Therefore, the missing Ahi1 domain is likely of the distal CC.

**Figure 4.**
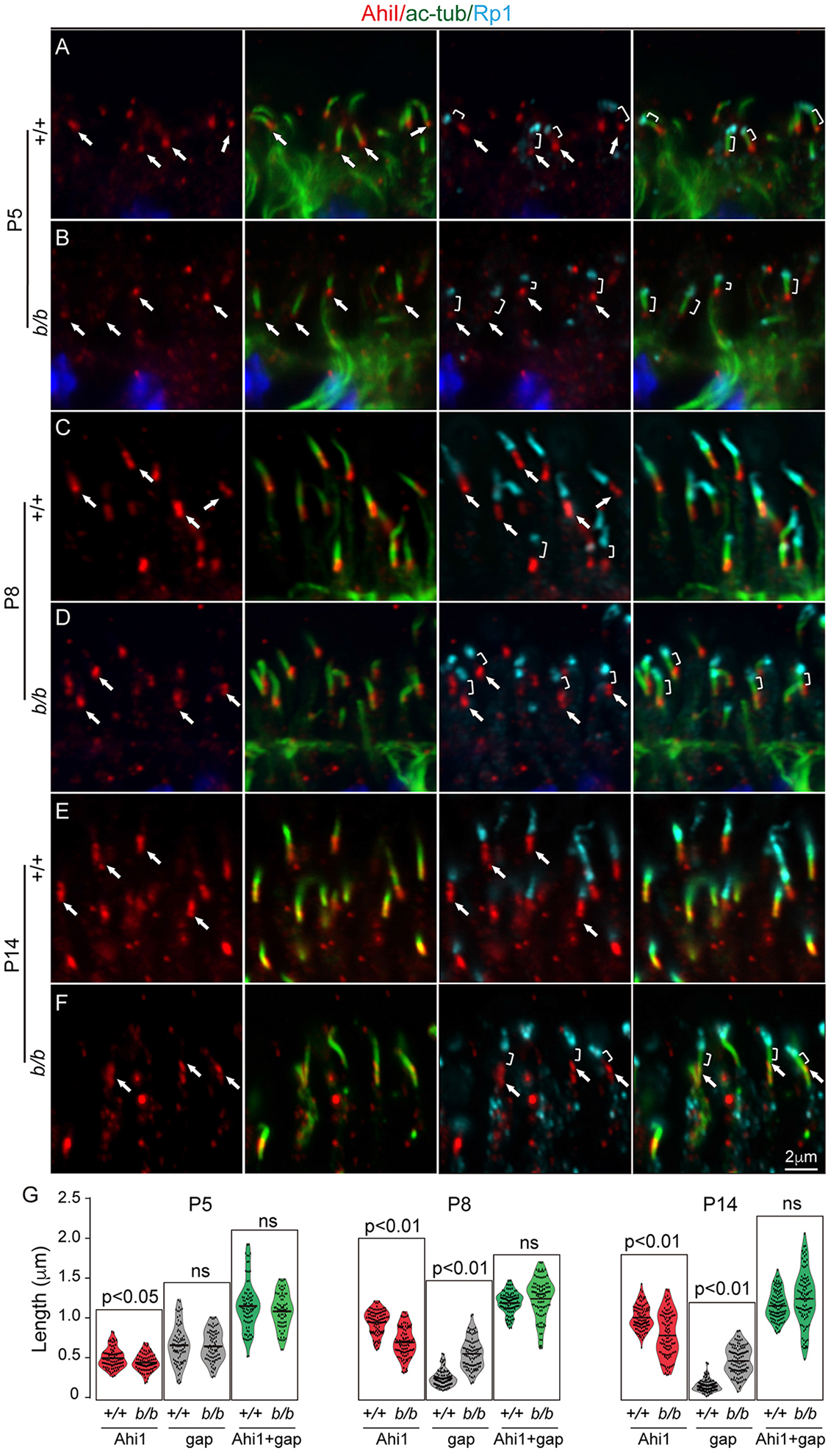
Mislocalization of connecting cilium protein Ahi1 in *Tmem138^b/b^* photoreceptors. Ahi1, Ac-tub, and Rp1 triple labeling for all panels. Red, green, and turquoise channels represent Ahi1, Ac-tub, and Rp1, respectively. (A-B), P5 sections. Arrows point to Ahi1 staining, while brackets indicate gaps between Ahi1 and Rp1 staining. (C-D), P8 sections. (E-F), P14 sections. (G), Quantification of Ahi1 domain length, gap length between Ahi1 and Rp1, and Ahi1 domain length plus gap length. The total domain length of “Ahi1+gap” is roughly equal between wild type and the mutants, suggesting the gap of the mutant distal CC is a missing part of Ahi1 domain.

We next examined several other ciliary proteins including Cep164, a protein in ciliary appendages (Graser et al., 2007), and IFT components Ift88 and Kif3a, for anterograde protein transport to the OS. The Cep164 staining appears to be identical in both the wild type and mutant photoreceptors through all ages examined (Fig. 5A-F, red). While appearing normal at P5 (Fig. 5G, H), the basal cilia staining of Ift88 was attenuated in P8 and P14 mutant photoreceptors with increased intensity of apical staining (Fig. 5I-L). In contrast Kif3a did not exhibit notable changes in the mutant even at age of P14 (Suppl. Fig. 3). Quantification of the domain length of acetylated α-tubulin did not show significant differences in contrast to Rp1, which was consistently shortened at P8 and P14 (Fig. 4, Fig.5 M-O). Thus, these data indicates that CC and axonemal compartments were severely perturbed upon disruption of *Tmem138*.

**Figure 5.**
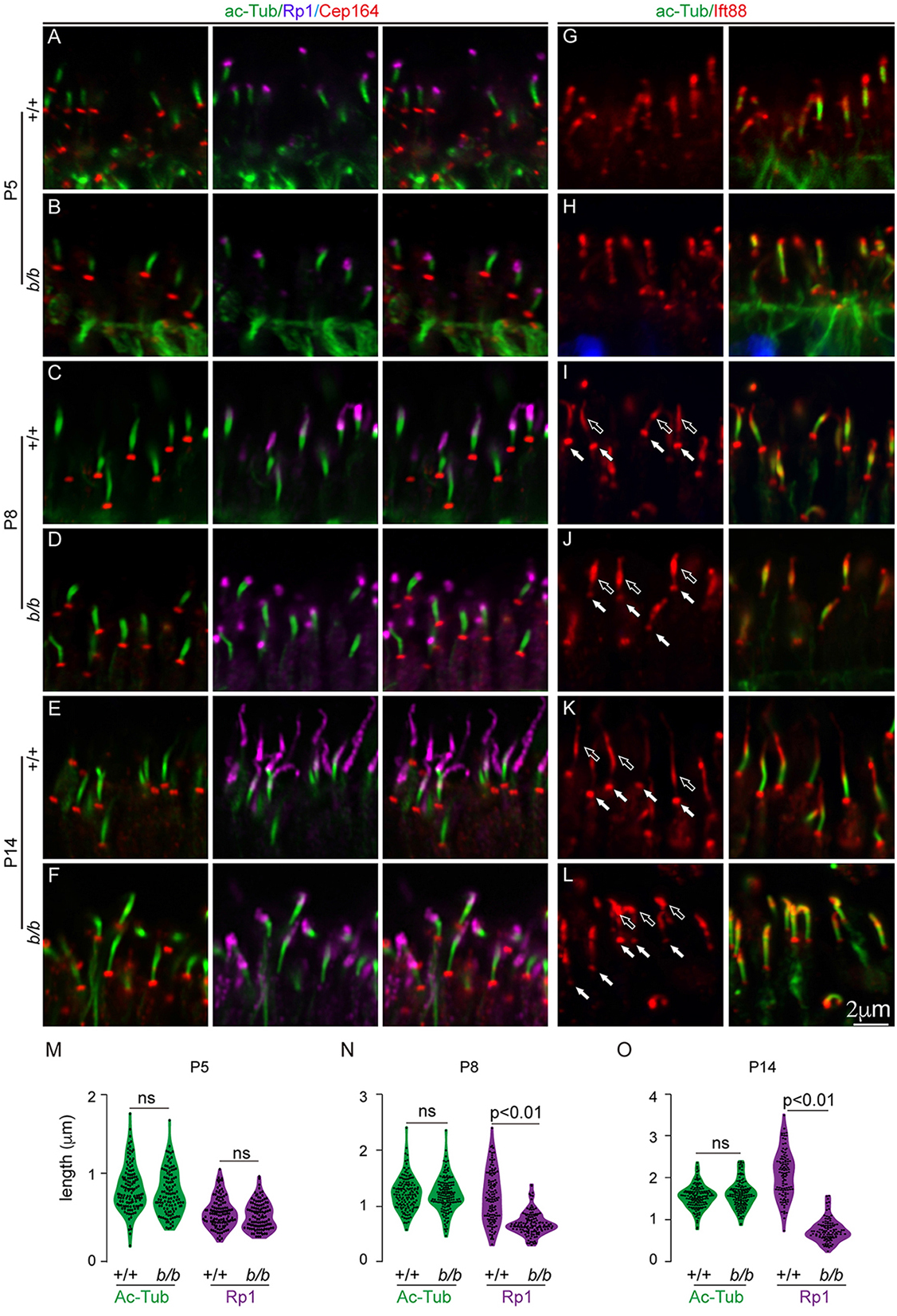
Altered ciliary domains in the mutant photoreceptor of *Tmem138^b/b^* retina. (A-F), Triple staining of ac-tubulin (spanning CC), Rp1(for axoneme), and Cep164 (for distal appendages). Colored protein names on top of the figure correspond to colors of imaging channels. All three proteins showed similar expression and localization in both wild type and the mutant cilia at P5. Shortened axonemes of the mutant cilia were observed at P8 and P14. (G-L), Double staining of ac-tubulin (for CC) and Ift88 (for CC and axonemes). Shortened axonemes in the mutants were observed at P14. (M-O), Quantification of CC (ac-tub staining) and axonemal (Rp1 staining) lengths. Student *t-test* was performed to detect statistical power. ns, no significance (p>0.01).

### Photoreceptor ciliogenesis appeared normal in the *Tmem138* mutants at ultrastructural levels

We next investigated whether ciliogenesis was affected by disruption of *Tmem138* using scanning electron microscopy (SEM) and TEM. SEM imaging demonstrated similar ciliary morphology of both wild type and mutant photoreceptor cells at P5 (Fig. 6A, B), when most photoreceptor cells did not develop OS. At P8 & P14, wild type photoreceptors began to grow OS distally from the CC (Fig. 6C, E), whereas no OS were found in the mutant cells (Fig. 6D, F). To obtain further details of photoreceptor ciliogenesis, we performed TEM to examine early developmental ages of photoreceptors. Photoreceptor ciliogenesis can be divided into 4 stages (Sedmak and Wolfrum, 2011): S1, ciliary vesicle docking on distal end of the mother centriole (Fig. 6G); S2, ciliary shaft elongation within the ciliary vesicle (Fig. 6H); S3, membrane fusion of the ciliary vesicle with the apical surface of IS (Fig. 6I); and S4, elongation of ciliary shaft to assemble axoneme (Fig. 6J). Multiple TEM sections collected from both wild type and mutant retinas showed comparable ciliary features at each of the four stages (Fig. 6G-J). Two additional stages S5 and S6, pertaining specifically to photoreceptor OS growing phase between P5 and P7 (Fig. 6K), were not found in mutant photoreceptors. In contrast, the array of 9 microtubule doublets of the mutant CC were similar to that of wild type photoreceptors (Fig. 6L). Furthermore, the frequency in finding each stage of S1∼S4 ciliogenesis was comparable between wild type and mutant photoreceptors (Fig. 6M).

**Figure 6.**
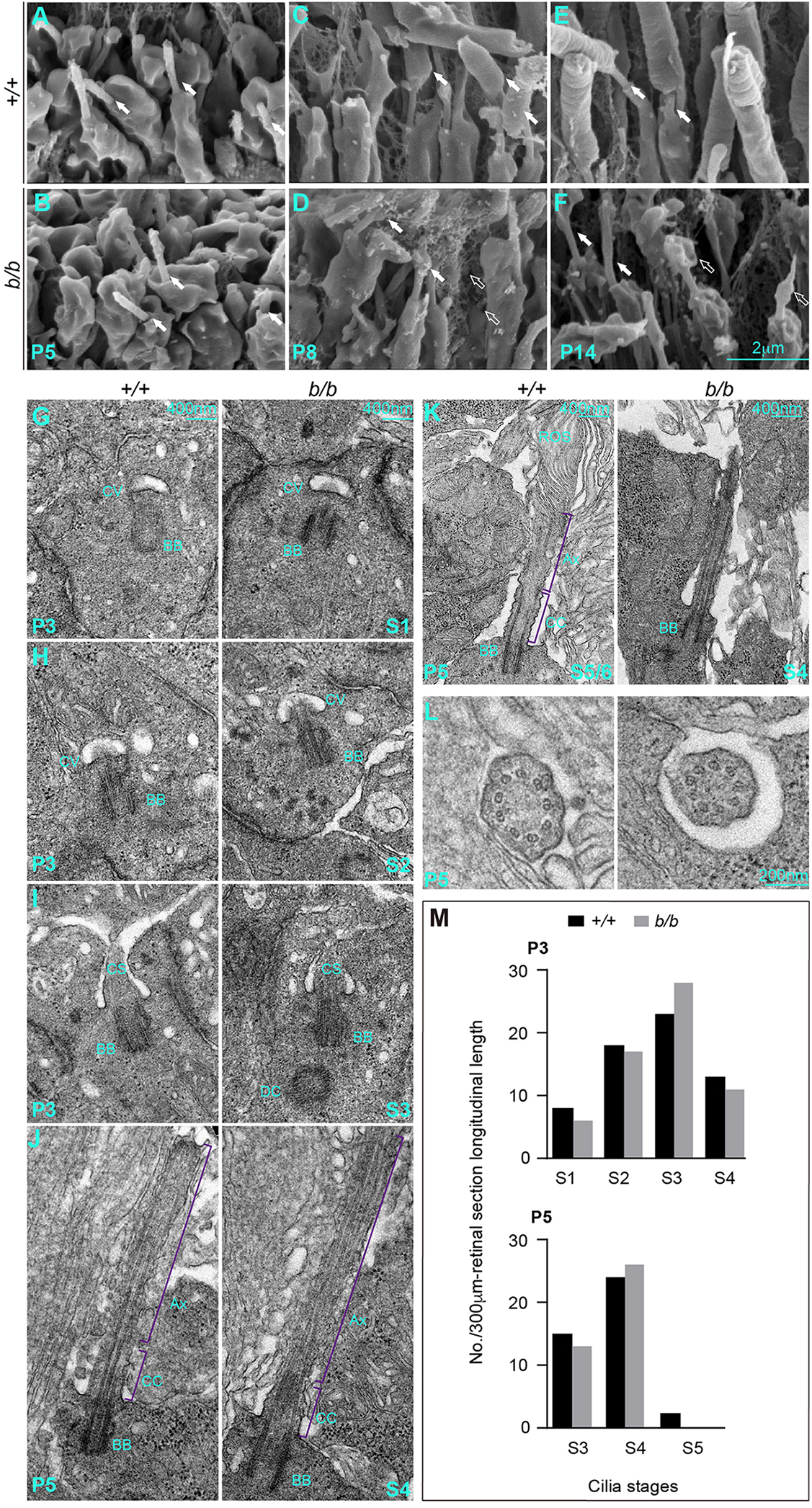
Photoreceptor ciliogenesis revealed by scanning electron microscopy (SEM) and transmission electron microscopy (TEM). (A, B), SEM pictures of wild type (A) and mutant (B) photoreceptor cilia (arrows) at P5. (C, D), P8 photoreceptors with arrows point to the OS in (C) and membrane stuff in (D). Open arrows point to the membrane vesicles embedded in the interphotoreceptor matrices. (E, F), Arrows point to the photoreceptor CC; open arrows point to the membranous materials on top of the mutant cilia. (G-J), P3 retinal sections. (G), Stage I cilium (S1): Ciliary vesicles docked on the distal end of the mother centrioles. (H), S2 cilium: Ciliary shafts elongated within the ciliary vesicles. (I), S3 cilium: Ciliary vesicle membrane fused with that of the IS at the apical surface. (J), S4 cilium: Cilium grew to assemble axoneme. (K, L), P5 retinal sections. (K), S5/S6: rudimental OS emerged above the cilia in wild type but not mutant photoreceptors. (L). Sections across the CC showed 9 doublet microtubules in both wild type and mutant photoreceptor cells. (M), Quantification of cilia numbers at each stage at P3 and P5. Cilia between RPE and the ONL at each stage of photoreceptor ciliogenesis were counted from 3 retinal sections/retina, each longitudinally spanning over a 100 μm-length in mid-peripheral retinal areas. The numbers of cilia from 3 sections (3×100 μm=300 μm total span) for each stage were used for plotting the graphs. One retina for each genotype was counted.

We further quantified numbers of cilia by using Ac-tub-stained flat mount retinas under light microscopy (Suppl. Fig. 4A-C). No significant changes in cilia numbers were detected at P8 between the wild type and mutant retinas (Suppl. Fig. 4D-F, J). Reduction in cilia numbers of the mutants at central and middle retinal areas at P14 (Suppl. Fig. 4G-I, K) probably reflects the previously observed photoreceptor death. Additionally, the general reduction in cilia density at P14 in comparison with that of P8 in both wild type and mutant photoreceptors was likely a result of retinal growth and expansion (Suppl. Fig. 4L). Taken together, the results of SEM, TEM, and light microscopy indicate that early stages of ciliogenesis of the mutant photoreceptors appeared structurally normal.

### Misrouting rhodopsin prior to initiation of the OS growth

To obtain functional insights of Tmem138 in OS biogenesis, we examined a series of photoreceptor developmental ages from P3 to P14 by monitoring the expression and localization of rhodopsin (Rho), the most abundant OS protein. At P3, as the inner segment (IS) starts to form, apical rhodopsin positive puncta were detected in both wild type and mutant retinas; rhodopsin was also evenly distributed in the inner segments and cell bodies (Suppl. Fig. 5A, B). At P4, the connecting cilium emerged apically in both wild type and mutant retinas (Suppl. Fig. 5C, D); However, rhodopsin was more intense at the wild type CC than that of the mutant, and a similar level of rhodopsin was seen in inner segments and cell bodies of both wild type and mutant photoreceptors (Suppl. Fig. 5C, D). Starting at P5, rhodopsin localization in the wild type cells was completely shifted to the ciliary protrusions, but remained randomly distributed throughout the mutant cell bodies (Fig. 7A, B). At P8 and P14, sprouting OS with strong and compartmentalized localization of rhodopsin and Pde6B were well developed in WT photoreceptors, whereas they appeared to be degenerating in the mutant with lower levels of rhodopsin and Pde6B that distributed randomly throughout the cell bodies (Fig. 7C-F).

**Figure 7.**
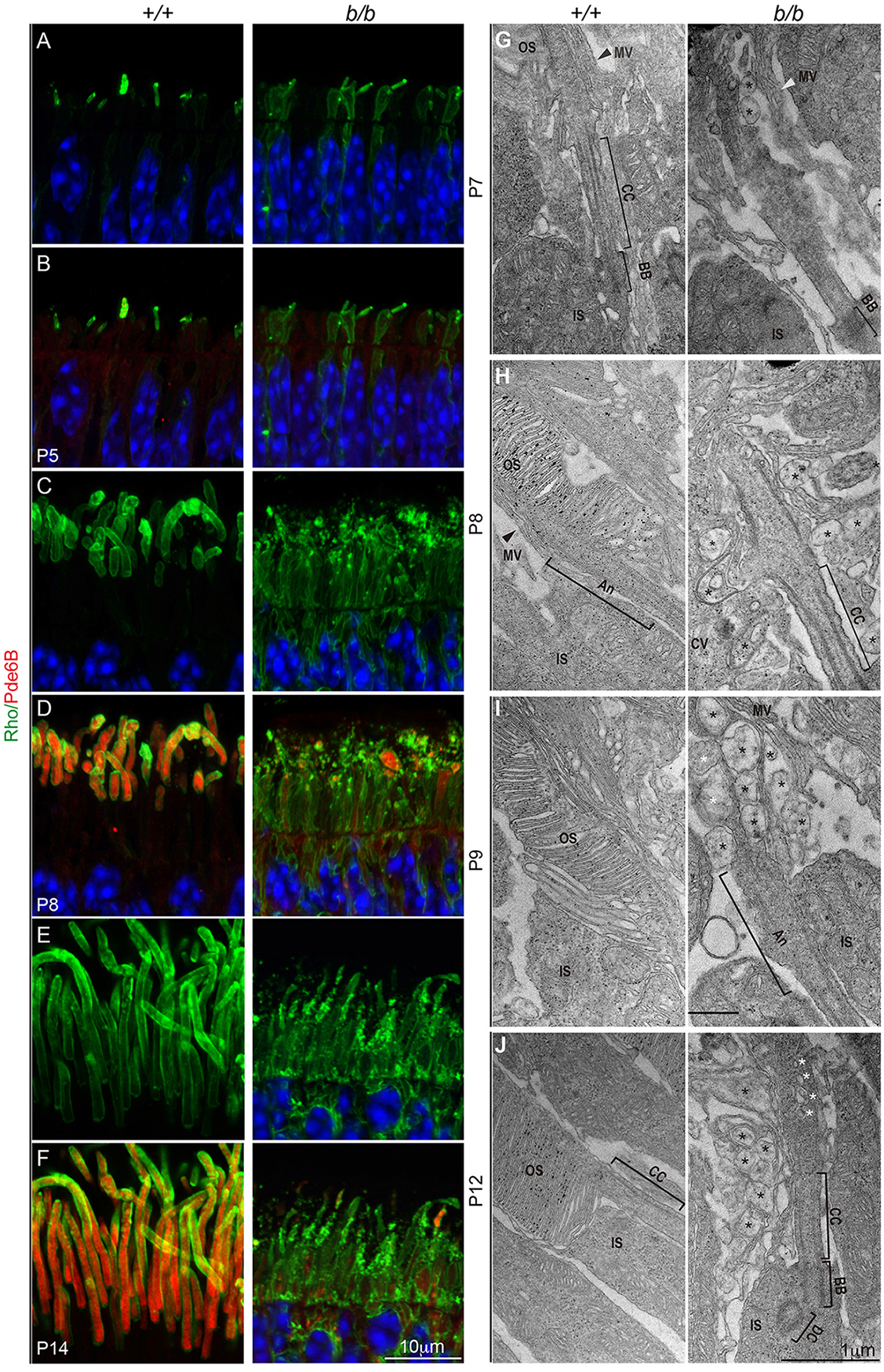
Failure of photoreceptor OS formation. (A-E) Photoreceptor OS was labeled with antibodies against Rho and Pde6B. (A, B), Mislocalization of rhodopsin was detected as early as P5 in the mutant retina. Note that Pde6B was only weakly expressed at P5. (C-F), At P8 and P14, Rho and Pde6B were strongly expressed and transported to the OS in the wild type photoreceptors but were mislocalized throughout the mutant photoreceptor cell body. Note rhodopsin positive puncta in the presumptive OS and IS regions and absence of intact OS in the mutant retina. (G-J) Transmission electron microscopy (TEM) observations of photoreceptor OS development. MV (arrowheads), microvilli of retinal pigmental epithelium (RPE); CC, connecting cilium; BB, basal body; An, axoneme; DC, daughter centriole; black asterisks, aberrant membrane-bound vacuoles; white asterisks, vesicles within the ciliary shaft.

On TEM sections, membranous vesicles of varying sizes accumulated extracellularly surrounding the connecting cilia of the mutant photoreceptors from P7 to P12 (Fig. 7G-J), while occasional intraciliary vesicles were also seen. The ciliary components of the mutant photoreceptor including the DC, BB, and CC appeared normal (Fig. 7J). On immunoelectron microscopy (immuno-EM) sections labeled with a rhodopsin antibody followed by a gold-conjugated secondary antibody, rudimentary OS was densely labeled with gold particles in P5 normal photoreceptors (Fig. 8A). However, disorganized membranous structures in place of OS were found in the mutant, that were weakly positive for rhodopsin, and strong rhodopsin labeling was found at the IS plasma membrane (Fig. 8A). Vesicular profiles of varying sizes were observed filling the extracellular space between photoreceptors, many of which were laden with gold particles (Fig. 8B, C). The EM results corroborate and expand the findings of light microscopy, which point to misrouting of rhodopsin and failure of OS formation at the very beginning of the OS morphogenesis.

**Figure 8.**
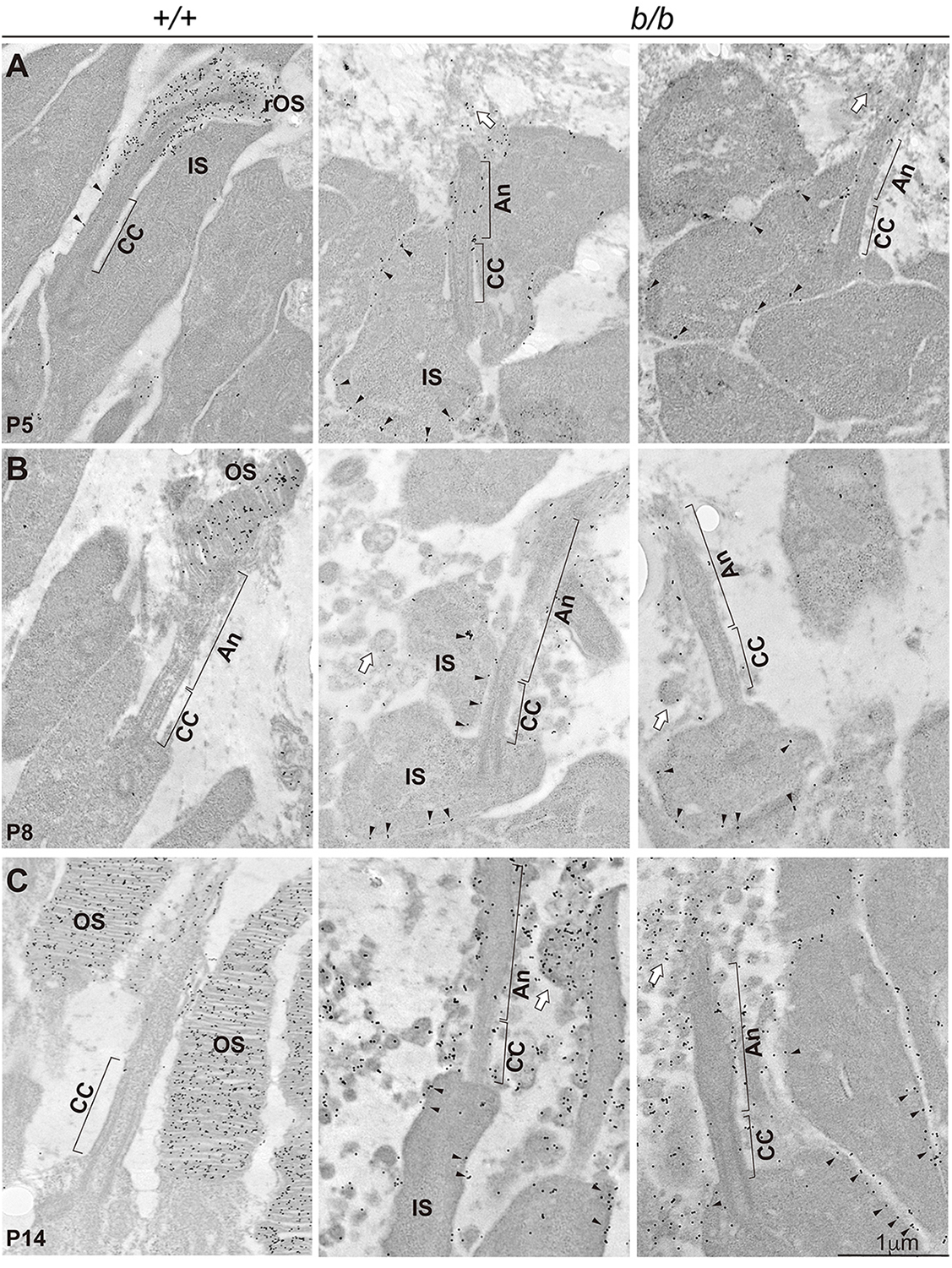
Impeded rhodopsin transport to the mutant OS revealed by immunoelectron microscopy (immuno-EM). (A). P5 retina sections. Immunogold particles-labeled rhodopsin was localized in the CC membrane and rudimentary OS (rOS) in the wild type photoreceptors but mislocalized to the IS membrane in the mutant. Some rhodopsin labels were observed near the distal end of the CC, probably associated with the failed OS morphogenesis. Black arrowheads point to rhodopsin labeling of the connecting cilium in the wild type photoreceptors but of the IS membrane in the mutant. Open arrows point to the disorganized membrane. (B), P8 retina sections. Disc structures appeared in the wild type OS packed with rhodopsin. Mutant photoreceptors were much like that of P5, except for the appearance of extracellular vesicles, some of which were rhodopsin positive. (C), P14 sections. Wild type OS was well developed, whereas mutant photoreceptors accumulated numerous rhodopsin-bearing extracellular vesicles.

### Continued functional requirement for Tmem138 in photoreceptor homeostasis

We next investigated whether Tmem138 was also crucial for OS maintenance once it had formed. *Actin-Flp* transgenic line was crossed onto *Tmem138^a/+^* mouse to generate a conditional knockout allele (*Tmem138^c^*), after which *Rhodopsin-Cre* line was used to delete E2 and E3 of the gene (Suppl. Fig. 6A, B). To increase excision efficiency, we produced *Rhodopsin-Cre;Tmem138^b/c^* compound mutants with one conditional allele (*Tmem138^c^*) on top of the germline null allele (*Tmem138^b^*). We first examined the timing of Cre expression and deletion efficacy of *Tmem138* by performing IHC using antibodies against Cre and Tmem138. Rhodopsin promoter-driven Cre expression was only detectable in a few cells at P8, more at P10 (Suppl. Fig. 6C, D), and widespread expression was observed between P14 and P21 (Suppl. Fig. 6D). Accordingly, Tmem138 protein was nearly completely ablated after P14 (Suppl. Fig. 6D). Rhodopsin-stained OSs were disorganized in many mutant photoreceptors at P14, with increased rhodopsin puncta near the base of the OS (Fig. 9A, B). Disorganization of the photoreceptors deteriorated further in 2-month-old mutant retinas (Fig. 9C, D). We next investigated whether ciliary compartments or protein transporting machinery were altered as they did in the germline knockouts. Similar to that of the germline knockouts, diminished and shortened Ahi1 and Rp1 domains (Fig. 9E-H), apical axonemal accumulation of Ift88 (Suppl. Fig.7A, B, E-H), and unaltered Kif3a (Suppl. Fig.7C, D) were observed, consistent with that of the germline knockouts.

**Figure 9.**
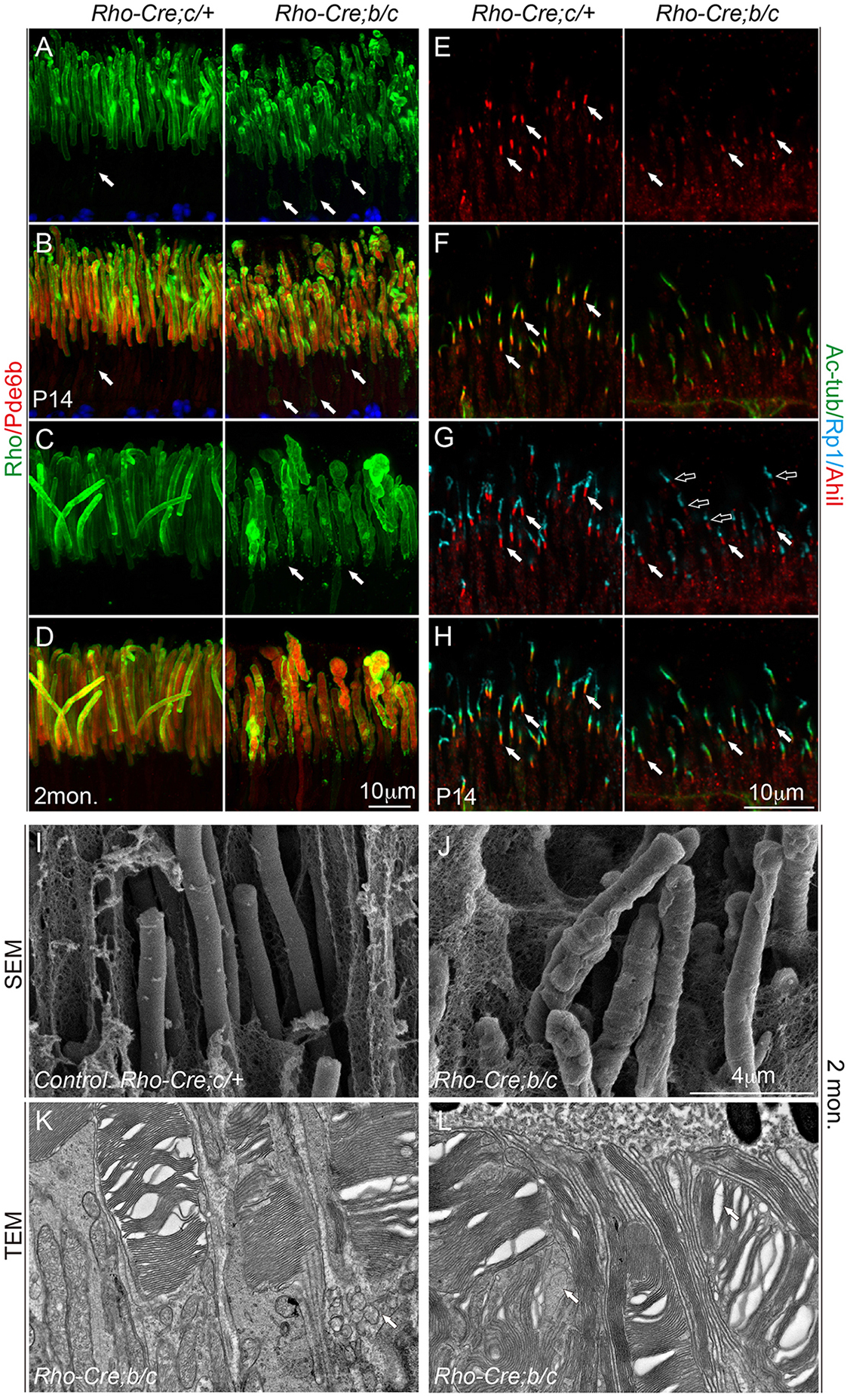
Photoreceptors of *Tmem138 ^b/c^* conditional knockout mice. (A, B), P14 photoreceptors. Disorganized OS and mislocalization of rhodopsin to the IS were revealed by Rho and Pde6b staining. (C, D), 2-month-old (2mon.) photoreceptors. The mutant photoreceptors appeared swellen. (E-H) P14 photoreceptors. Shortened and diminished Ahi1 and Rp1 staining was observed. (I, J), SEM pictures of wild type (I) and mutant (J) photoreceptors. (K, L), Two representative images of mutant photoreceptors showing dysmorphic OS. Arrows point to inter-(K) and intra-(L) photoreceptor OS vesicles.

At ultrastructural levels, 2-month-old mutant photoreceptors exhibited bumpy outer surfaces in comparison with the smooth and uniform OS in the wild type on SEM sections (Fig. 9I, J). On TEM sections, rod OS discs showed variable morphological defects including splitting and disorientated discs, intra- and inter-photoreceptor membrane vacuoles (Fig. 9K, L). These results suggested that Tmem138 functions not only in the initial OS morphogenesis but also in OS renewal, likely through similar mechanisms as in the early phase of the OS development.

### Tmem138 interacts with rhodopsin, Ahi1 and Tmem231

We next explored possible mechanisms by whichTmem138 facilitates rhodopsin transport. We first tested whether Tmem138 directly interacts with rhodopsin by a protein pulldown assay. We co-expressed 3xFlag-TST (Twin-Strep-tag)-tagged Tmem138 and HA-tagged rhodopsin in HEK 293 cells, and purified the protein extracts through columns loaded with Strep-Tactin® resins and HA antibody cross-linked Sepharose beads (see Materials and Methods). In either forward or reverse direction, we were able to detect an interaction between Tmem138 and rhodopsin (Fig. 10A). Furthermore, Rhodopsin co-immunoprecipitated with Tmeme138 from the wild type retinal extracts but little rhodopsin was pulled down in the negative control (Fig. 10B), which was the conditional mutant retina with *Rhodopsin* promoter-driven *Cre* (*Rho-Cre)* (see later and Materials and Methods).

**Figure 10.**
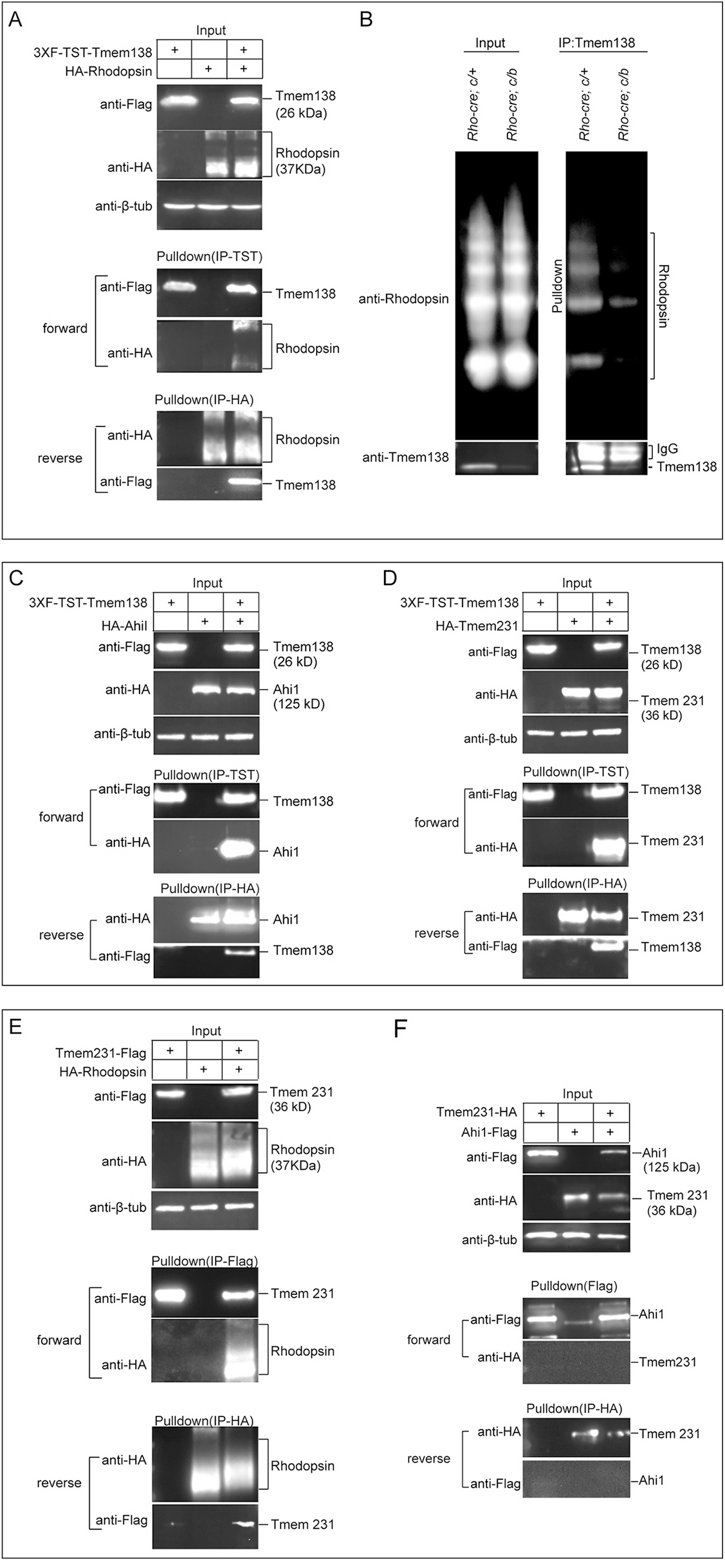
Tmem138 interacts with rhodopsin and JBTS proteins Ahi1 and Tmem231. (A), Tmem138 interacts with rhodopsin revealed by protein pull-down assay in HEK293 cells in both forward and reverse directions. (B), Tmem138 coimmunoprecipitated with rhodopsin from P21 retinal extracts. The control retinal extracts were prepared from *Rohdopsin-Cre*-driven *Tmem138 ^b/c^* conditional knockout retinas (See Suppl. Fig.6). (C), Tmem138 interacted with Ahi1. (D). Tmem138 interacted with Tmem231. (E) Tmem231 interacted with rhodopsin. (F), Tmeme231 did not interact with Ahi1.

A literature search found two JBTS genes, *Ahi1* and *Arl13b*, that when mutated in mice also lead to failed OS growth (Dilan et al., 2019; Hanke-Gogokhia et al., 2017; Louie et al., 2010). Another TZ localized JBTS/MKS protein Tmem231 has not been tested in photoreceptors, but mutations in mice caused microphthalmia during embryogenesis (Roberson et al., 2015; Srour et al., 2012). Using similar affinity columns described above, we found both Ahi1 and Tmem231 interacted with Tmem138 (Fig. 10C, D). Moreover, Tmem231 but not Ahi1 interacted with rhodopsin (Fig. 10E, Suppl. Fig. 8A), and the two did not interact with each other (Fig. 10F). We did not observe Arl13b interacting with Tmem138 (Suppl. 8B). Furthermore, neither Tmem67/Meckelin, which when mutated in mice blocked OS biogenesis (Collin et al., 2012), nor Rab8a, a GTPase important for rhodopsin transport from Golgi to the base of the CC, interacted with Tmem138 c(Suppl. Fig. 8C, D).

Because Tmem231 interacted with both Tmem138 and rhodopsin, we asked whether its localization was also changed in the mutant photoreceptors. Tmem231 appeared to be localized to a broad area of photoreceptor cilium including the base, CC and proximal axoneme (Suppl. Fig. 9A, B, E, F, I, J, M, N, Q, R). Tmem231staining was slightly weaker in the mutant CC compared with that of wild type photoreceptors at the examined ages (Suppl. Fig. 9B, D, F, H, N, P). Together, these results suggest that Tmem138 may recruit a subset of JBTS proteins directly involved rhodopsin transport across the cilium.

## Discussion

Despite extensive investigations, molecular processes of the photoreceptor OS biogenesis and related protein trafficking remain largely unresolved. To obtain further mechanistic insights into these processes and related disease mechanisms, we conducted the current study focusing on Tmem138, a putative transmembrane protein implicated in JBTS (Suzuki et al., 2016; Tuz et al., 2013) (Lee et al., 2012) with associated retinal dystrophy.

Inactivation of *Tmem138* in mice led to ciliopathy like phenotypes, consistent with previous reports of Tmem138 as a ciliary protein in worm and cultured mammalian cells (Jensen et al., 2015; Lee et al., 2012; Li et al., 2016). One of the most striking phenotypes in the *Tmem138* mutants is that photoreceptors completely fail to develop OS and undergo early-onset degeneration. To understand the role of Tmem138 in OS biogenesis, we first examined its protein localization in photoreceptors. Although it has been reported that Tmem138 is located at the base and axoneme of the cilium in cultured mouse IMCD3 cells (Lee et al., 2012), and in the TZ (equivalent to the photoreceptor CC) of *C. elegans* sensory neurons (Jensen et al., 2015; Li et al., 2016), its localization in photoreceptors had not been examined previously. Our finding that Tmem138 is localized to the CC of the photoreceptor is consistent with it being a TZ protein in the *C, elegans* and a ciliary protein in cultured cells. Futhermore, disruption of *Tmem138* altered ciliary compartments as evidenced by shortened domain of Ahi1 and Rp1, resident proteins in CC and axoneme respectively, suggesting Tmem138 functions primarily in the CC.

Several other cilia gene mutants do not form OS including *Cep290* (Rachel et al., 2015), a gene mutated in diverse ciliopathies (Coppieters et al., 2010) and two other JBTS genes, *Arl13b* and *Ahi1* (Dilan et al., 2019; Hanke-Gogokhia et al., 2017; Louie et al., 2010). *Cep290* mutant photoreceptors do not even form CC, which by itself is sufficient to abolish OS biogenesis (Rachel et al., 2015). The structural defects of photoreceptor cilia have not been closely examined in either *Arl13b* or *Ahi1* mutant mice (Dilan et al., 2019; Louie et al., 2010). In this study, we carefully examined photoreceptor ciliogenesis in the *Tmem138* mutant before cilia started to grow OS. The structural features of ciliogenesis at different stages (Sedmak and Wolfrum, 2011), the number of cilia, and the length of the CC all appeared normal before P8. However, mislocalization of rhodopsin was detected as early as P5. And Tmem138 interacted with rhodopsin in either cultured cell extracts with transfected constructs using protein pulldown assays or retinal extracts using co-immunoprecipitation. These results support the notion that Tmem138 functions in protein trafficking across the CC but probably not in ciliogenesis.

Genetic studies from *C. elegans* suggest TZ localization of Tmem138 depends on Cep290 and Mks5 (Rpgripl1) (Jensen et al., 2015; Li et al., 2016). TZ assembly pathways or interactors downstream of Tmem138, however, remains unknown, particularly for the photoreceptors. A search for candidate proteins interacting with Tmem138 identified rhodopsin, Ahi1/Juberin and Tmem231. The latter two are also JBTS proteins and localize to the CC compartment. Localizations of the two proteins are also altered in *Tmem138* mutant photoreceptors. Moreover, *Ahi1* mutant photoreceptors have similar phenotypes as the *Tmem138* mutants, as aforementioned, indicating that the Ahi1 and Tmem138 interact *in vivo*.

Tmem231 genetically interacts with multiple MKS proteins (Roberson et al., 2015). *Tmem231* mutations are associated with OFD6 syndrome, and so does Tmem138 (Li et al., 2016). The protein has a broad range of functions during early development. *Tmem231* mutant mice develop microphthalmia during early embryogenesis involving Shh signaling (Chih et al., 2011; Roberson et al., 2015). The protein is localized to the base, CC and proximal region of the axoneme of the photoreceptor cilia, and its CC staining is diminished in *Tmem138* mutants. Interestingly, both Tmem138 and Tmem231 directly interact with rhodopsin, and all three are membrane proteins. The interaction between CC membrane proteins with rhodopsin must be transient and dynamic, as the latter is in rapid transit through the CC on its way to the nascent OS discs. Tmem138 and Tmem231 do not seem to travel with rhodopsin, or at least not in one-to-one molar ratio. Otherwise they would be detected in the OS. Beside rhodopsin, Tmem138 likely mediates transport of other proteins to the OS since ablation of *rhodopsin* gene alone lead to a phenotype that is much less severe than that of *Tmem138* mutant (Lem et al., 1999). Nevertheless, we found that it is not feasible to examine other OS proteins at early age of OS biogenesis due to their much lower expression compared with that of rhodopsin, for example, the Pde6B.

Although Alr13b and Tmem138 have different ciliary localization in photoreceptors, the failure of OS biogenesis, the shortened Rp1 occupied domain and altered Ift88 distribution are common features in the mutant mice of both genes. The undetectable protein interaction between Arl13b and Tmem138 suggests that the two proteins might act at different steps during OS morphogenesis. We noted that the mouse mutant of another TZ membrane protein, *Tmem67/Meckelin*, develops a severe photoreceptor degeneration and lacks OS, but undergoes normal ciliogenesis (Collin et al., 2012). However, Tmem67 does not interact with Tmem138. Further testing of GTPases Rab8a, crucial for rhodopsin endomembrane protein trafficking of the photoreceptor (Wang and Deretic, 2014), did not result in their detectable interactions with Tmem138 either. The evidence taken together suggests that the Tmem138 functional domain is confined to the CC.

Finally, all molecular changes in the photoreceptor CC of the *Tmem138* germline knockouts are recapitulated in *Tmem138* conditional mutants having well-formed outer segments by the time of gene deletion, suggesting that Tmem138 is not only required for the initiation of OS biogenesis but also for its homeostasis using similary molecular machinery. We propose that Tmem138, Tmem231 and/or other CC proteins may form distinct complexes serving as guiding posts on the CC membrane for OS proteins (for example, rhodopsin) being transported across the CC membrane through transient interactions. Meanwhile, cytoplasmic interaction of Tmem138 with Ahi1 might serve as an anchor for other components to maintain CC integrity. Further experiments are apparently required to test such a model in the future.

## Acknowledgements

We thank Dr. Tiansen Li from the National Eye Institute for providing valuable suggestions in preparation of the manuscript. The authors thank Tiansen Li, Rong Ju for critical reading the manuscript and helpful comments. This work was supported by grants from the National Natural Science Foundation of China (NSFC: 31571077; Beijing, China), the Guangzhou City Sciences and Technologies Innovation Project (201707020009; Guangzhou, Guangdong Province, China), ‘‘100 People Plan’’ from Sun Yat-sen University (8300-18821104; Guangzhou, Guangdong Province, China), and research funding from the State Key Laboratory of Ophthalmology at Zhongshan Ophthalmic Center (303060202400339; Guangzhou, Guangdong Province, China) to Chunqiao Liu.

## Supplementary Figure legends

**Supplemental Figure 1.**
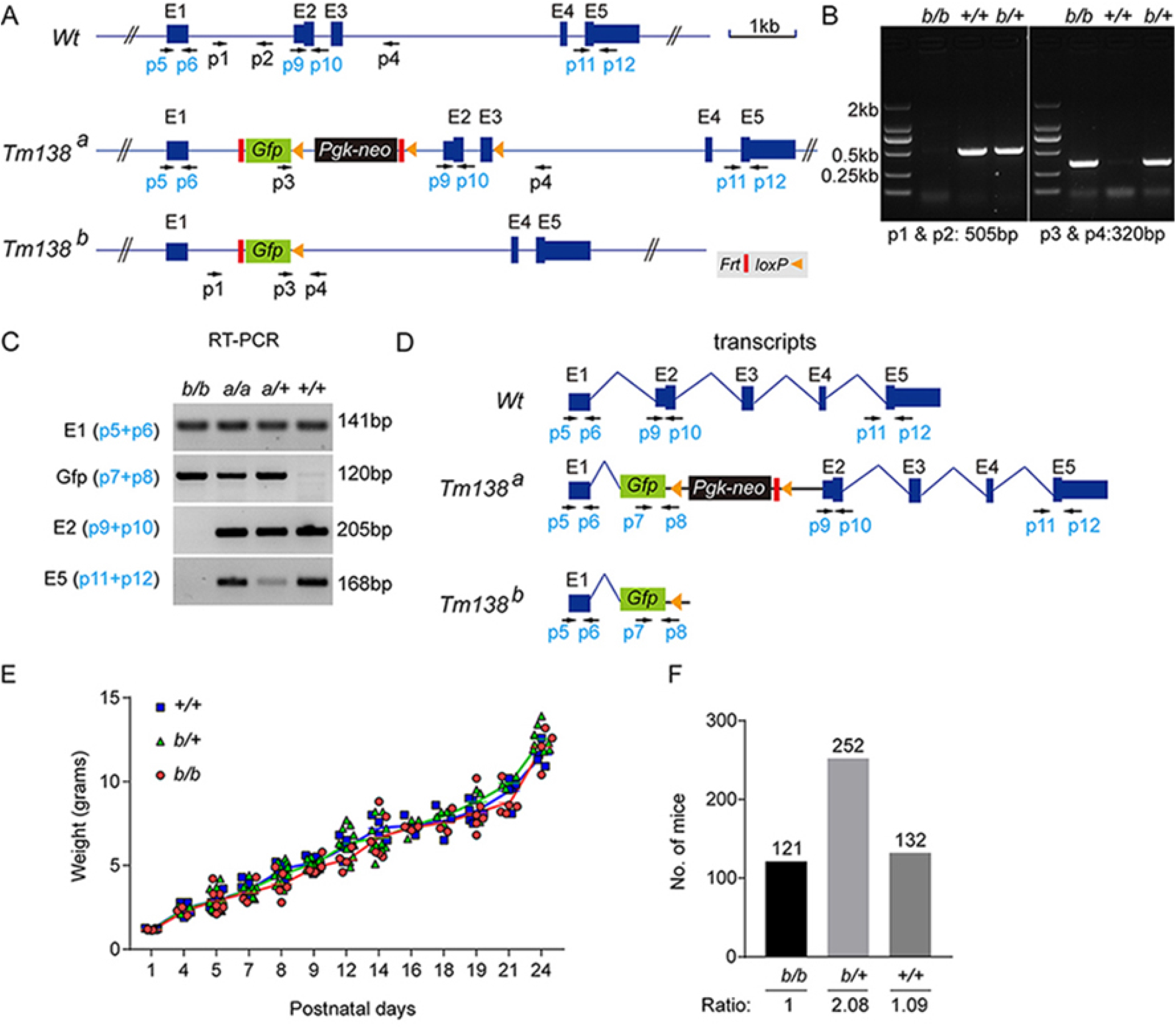
(A), Positions of primers used to differentiate genetic alleles and detect transcripts from each allele by RT-PCR. Primers in black (P1-P4) were used for genotyping. Primers in turquoise (P5-P12) were used for RT-qPCR. (B), Genotyping *Tmem138^+/+^*(*Wt, +/+*), *Tmem138^b/+^(b/+)*, and *Tmem138^b/b^(b/b)* mice. (C), RT-PCR-detected transcripts from selected exons. (D), Possible longest transcripts for each allele inferred from the results of (C). (E), Comparable growth curves from different genotypes. (F), Genotype ratio of pooled born mice.

**Supplemental Figure 2.**
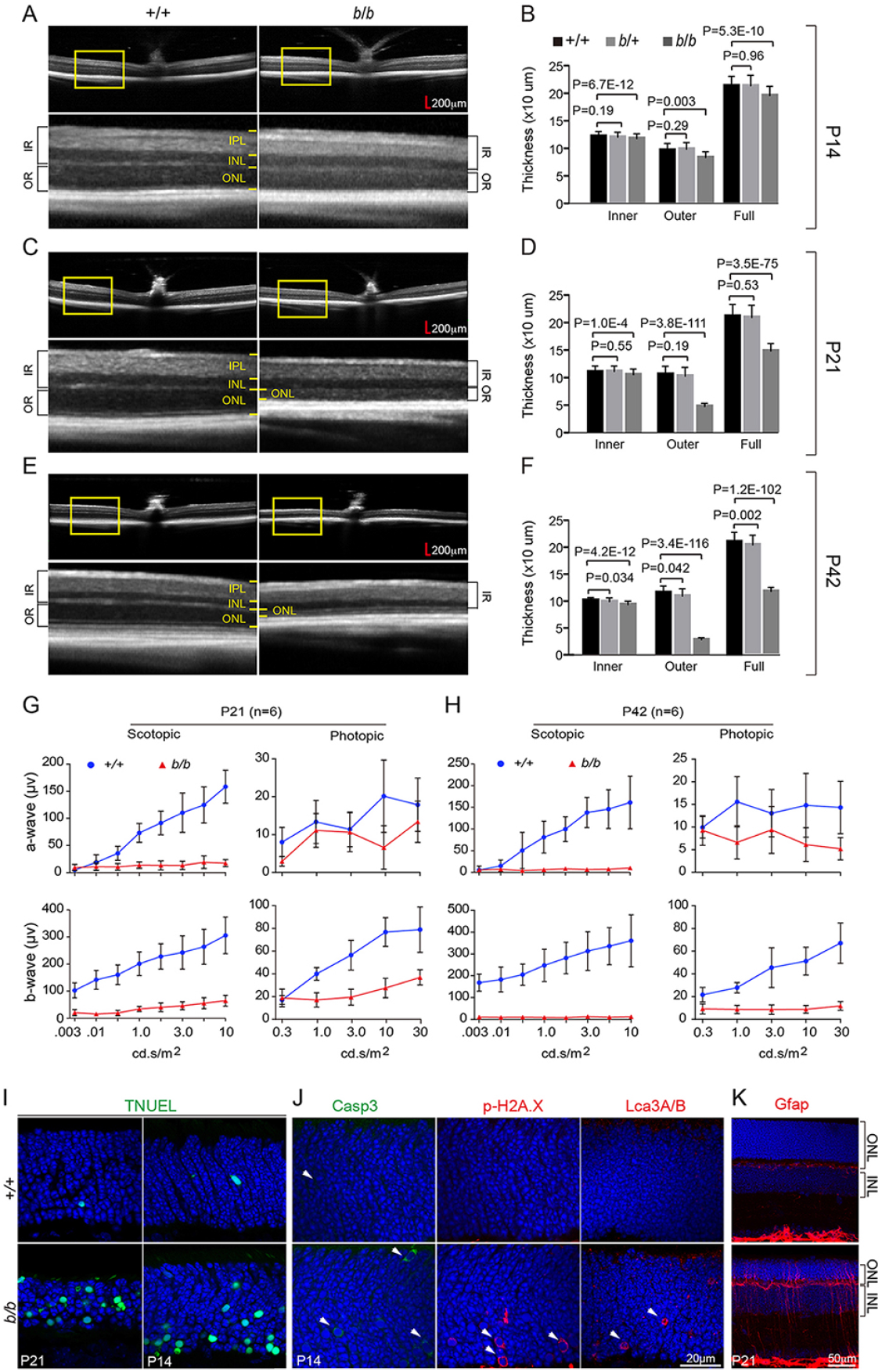
(A, B), (C, D), and (E, F) are OCT images and respective quantifications at P14, P21, and P42. Boxed areas were magnified below each panel. (G, H), Quantification scotopic and photopic a- and b-waves of ERG at P21 (G) and P42 (H). (I), Detection of cell death by TNUEL at P21 and P14. (J), Detection of cell death using anti-Caspase-3 and p-H2A. X antibodies, and detection of autophagy using anti-Lca3A/B antibody. (K), Muller glia activation revealed by GFAP staining.

**Supplemental Figure 3.**
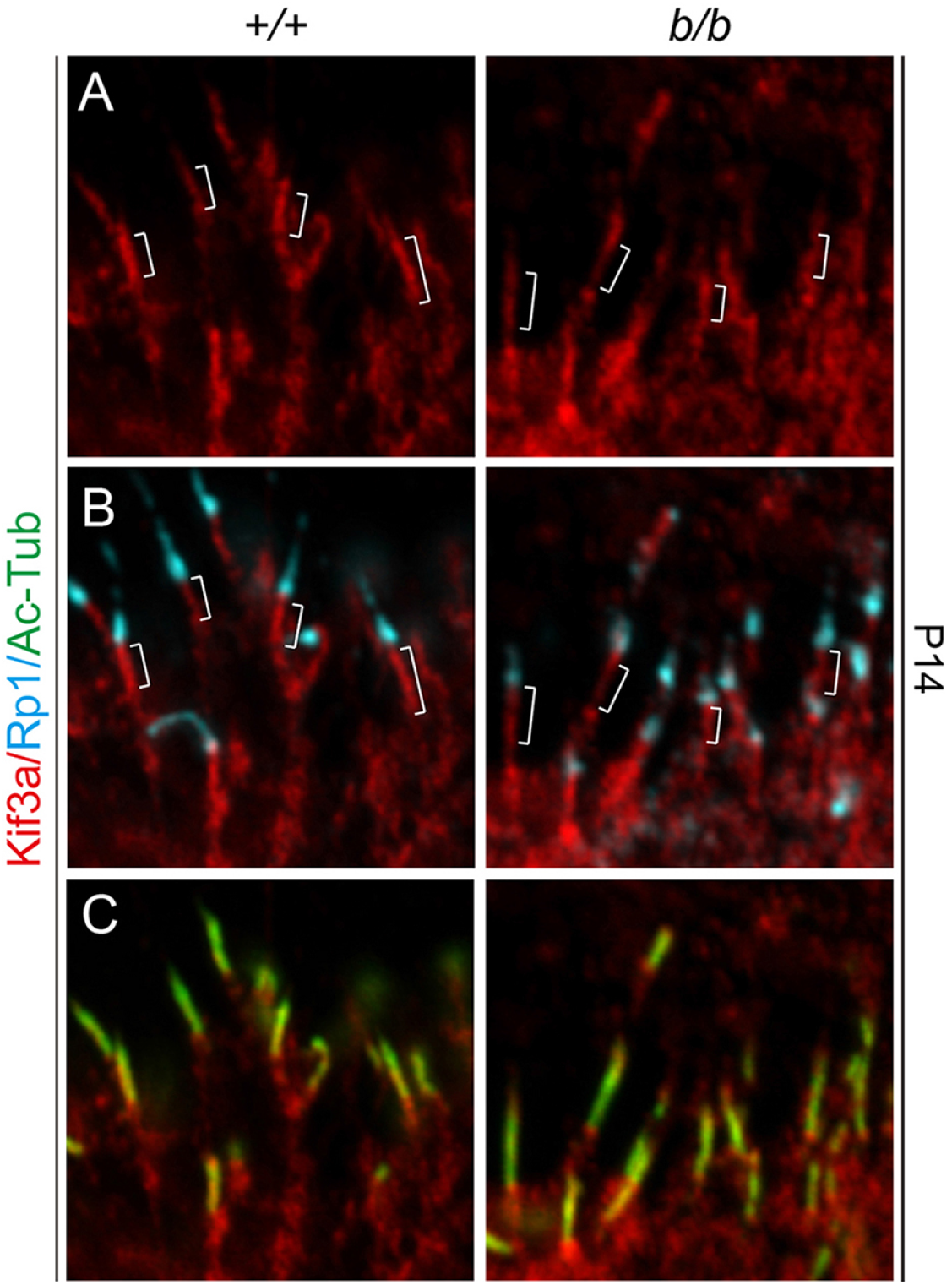
(A-C), Kif3a (red), Ac-tub (green), and Rp1(turquoise) triple labeling at P14. Brackets indicate Kif3a CC staining.

**Supplemental Figure 4.**
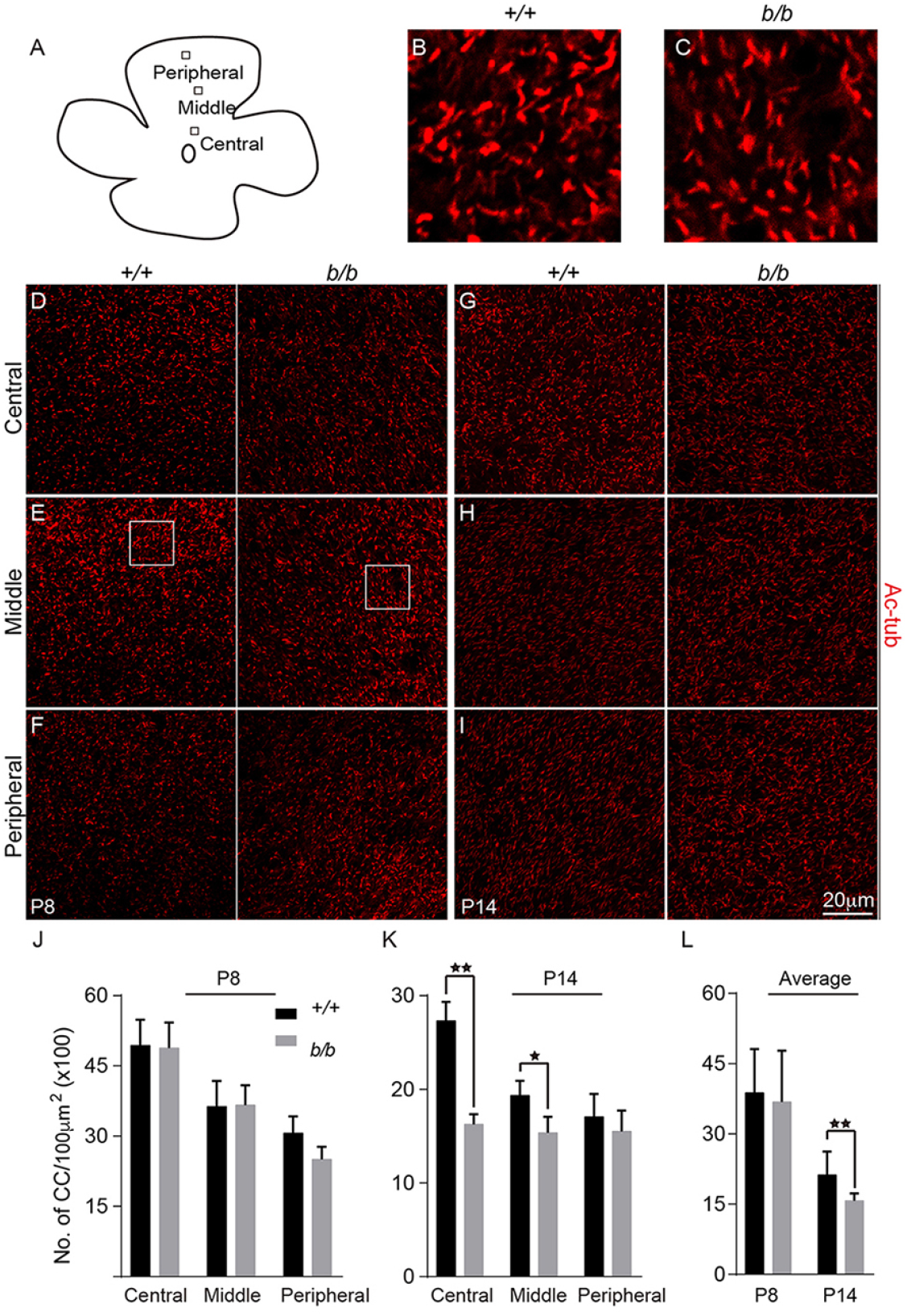
Quantification of cilia number by Ac-tub-stained flat mount retinas. (A), Schematic drawing of a retina with small squares indicating counted areas. (B, C), Magnified areas in (E) showing image examples for quantification of cilia numbers. (D-F), P8 retina with Ac-tub staining. (G-I), P14 retina with Ac-tub staining. (J, K), Cilia numbers of P8 (J) and P14 (K) counted retinal areas. (L), Average numbers of cilia from different areas from P8 and P14 retinas.

**Supplemental Figure 5.**
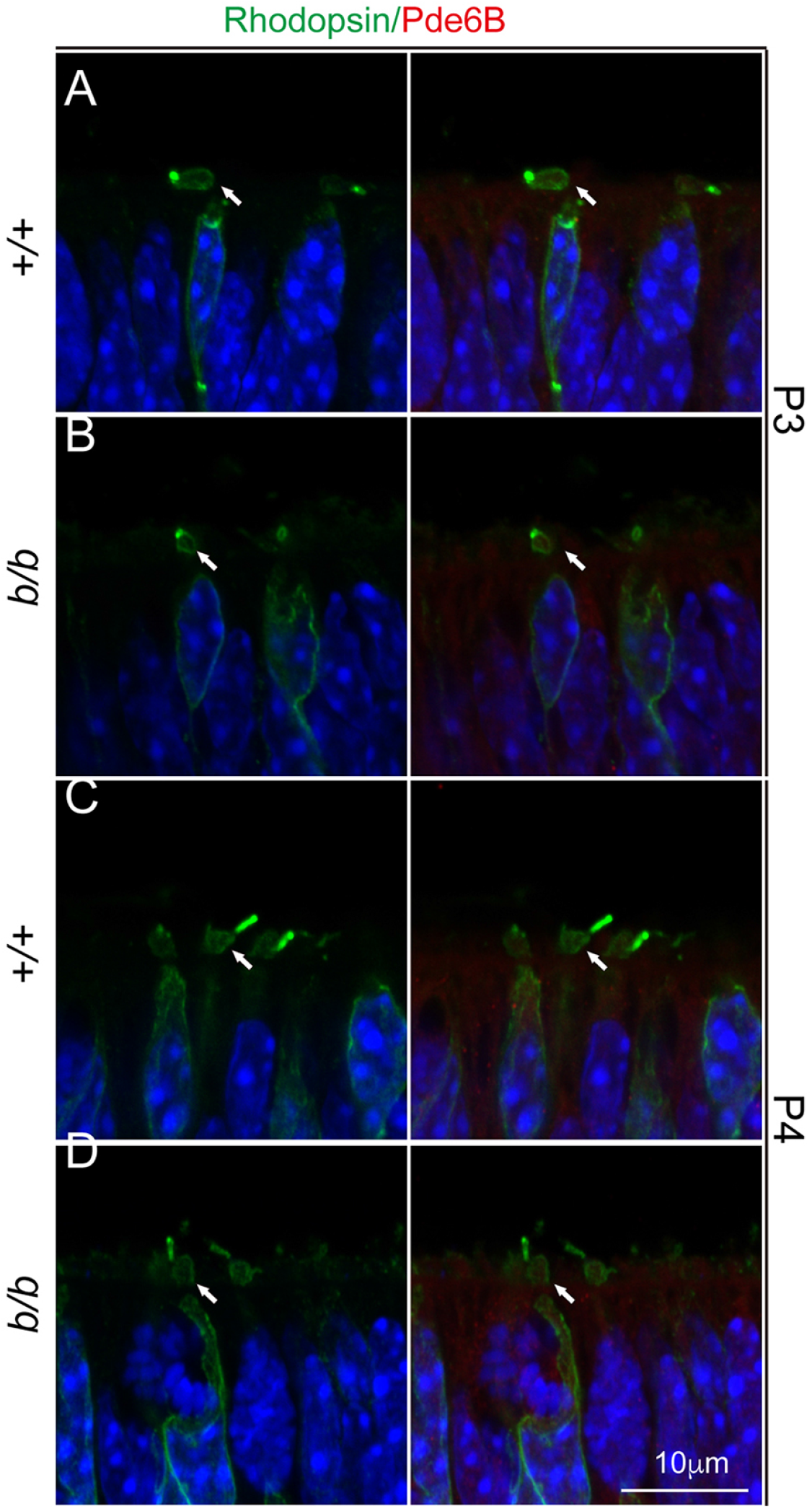
Rhodopsin- and Pde6B-stained P3 (A, B) and P4 (C, D) photoreceptor cells. Arrows point to the nascent inner segments.

**Supplemental Figure 6.**
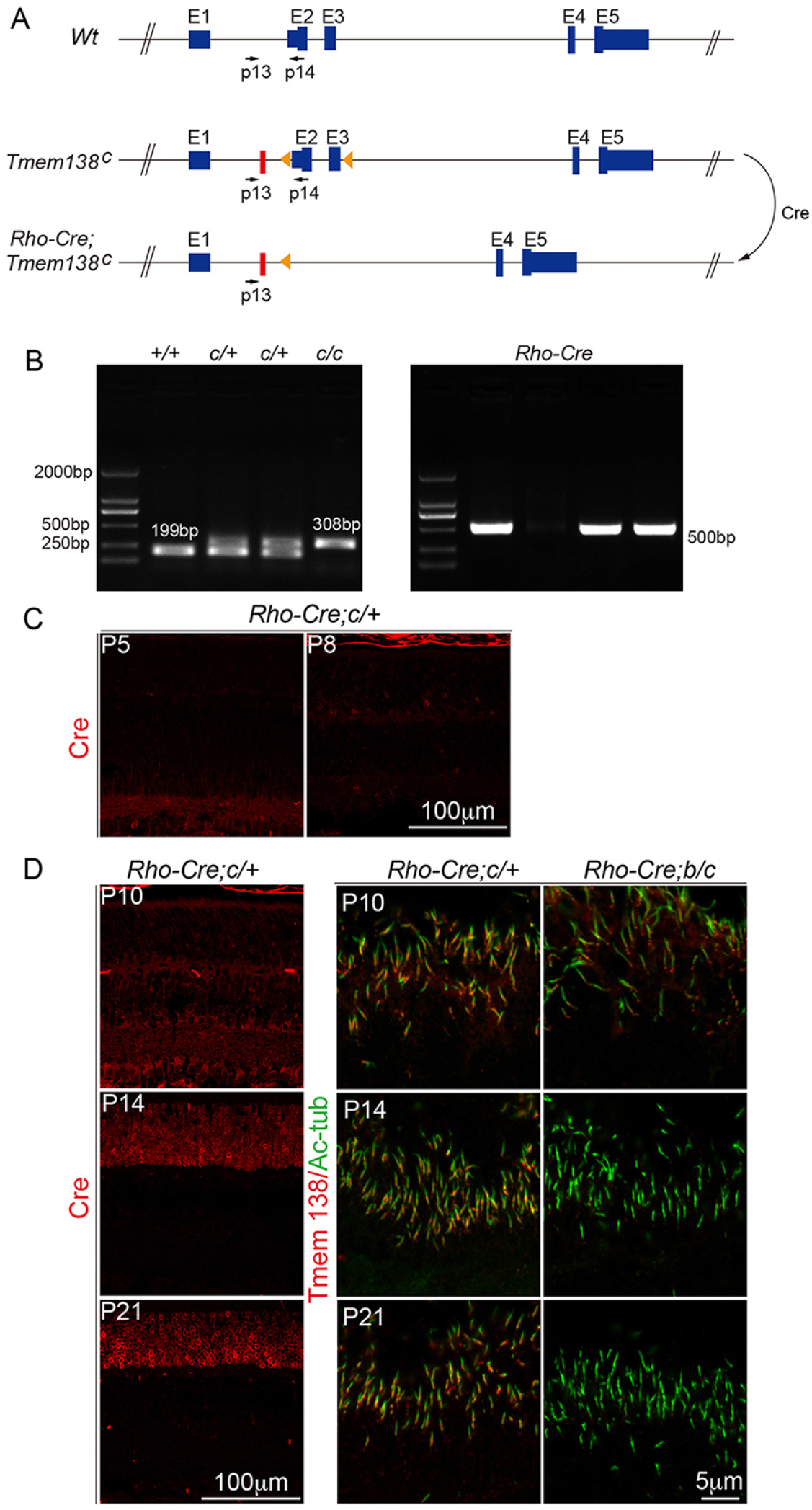
*Rhodopsin-Cre* (*Rho-Cre*) was used to conditionally ablate *Tmem138* from the *Tmem138^c^* allele (*Rhodopsin-Cre; c/+),* which was bred onto *Tmem138* null allele (*Tmem138^b/+^*) to create compound mutants *Rho-Cre;Tmem138^b/c^* mice (*Rhodopsin-Cre; b/c)*. (A) Locations of primers used for genotyping of *Tmem138^c/+^* (c/+) allele. (B), Left panel: genotyping of ‘c/+’ mice; right panel: genotyping of *Rho-Cre* mice; (C), Cre protein was not detected in P5, and sparsely detected in P8 retinas of *Rhodopsin-Cre; c/+* mice. (D), Cre was mildly expressed at P10 with attenuated Tmem138 expression. At P14 and P21, strong expression of Cre was detected in the ONL, and Tmem138 expression was almost gone.

**Supplemental Figure 7.**
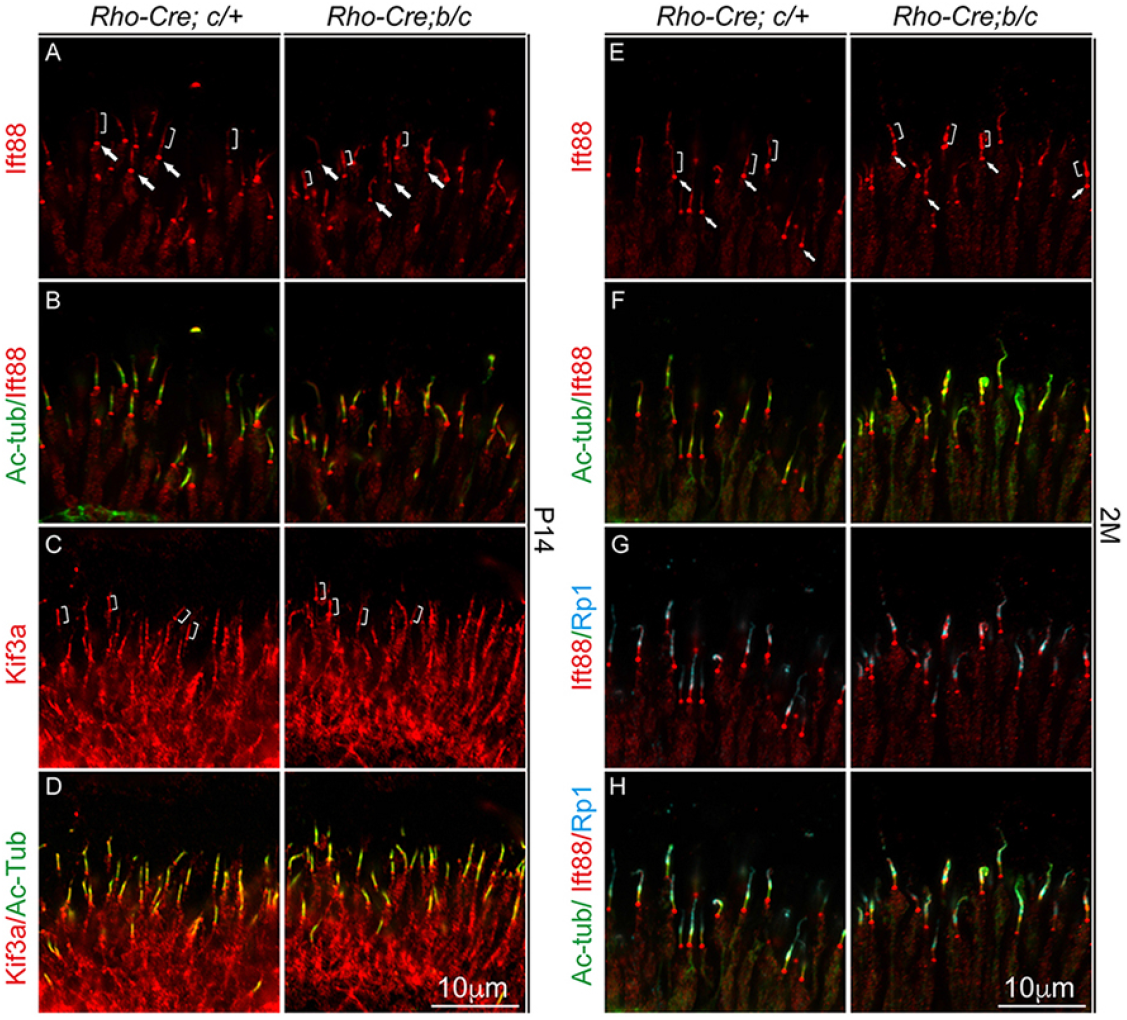
Genotypes of conditional mutants and controls were labeled on top of the figure. (A, B) Ift88 and Ac-tub labeled photoreceptor cilia at P14. (C, D) Kif3a and Ac-tub labeled P14 retina. (E-H) Ift88, Ac-tub and Rp1 triple-labeling photoreceptor cilia at P14. Arrows, base of the cilium; Brackets, proximal axoneme.

**Supplemental Figure 8.**
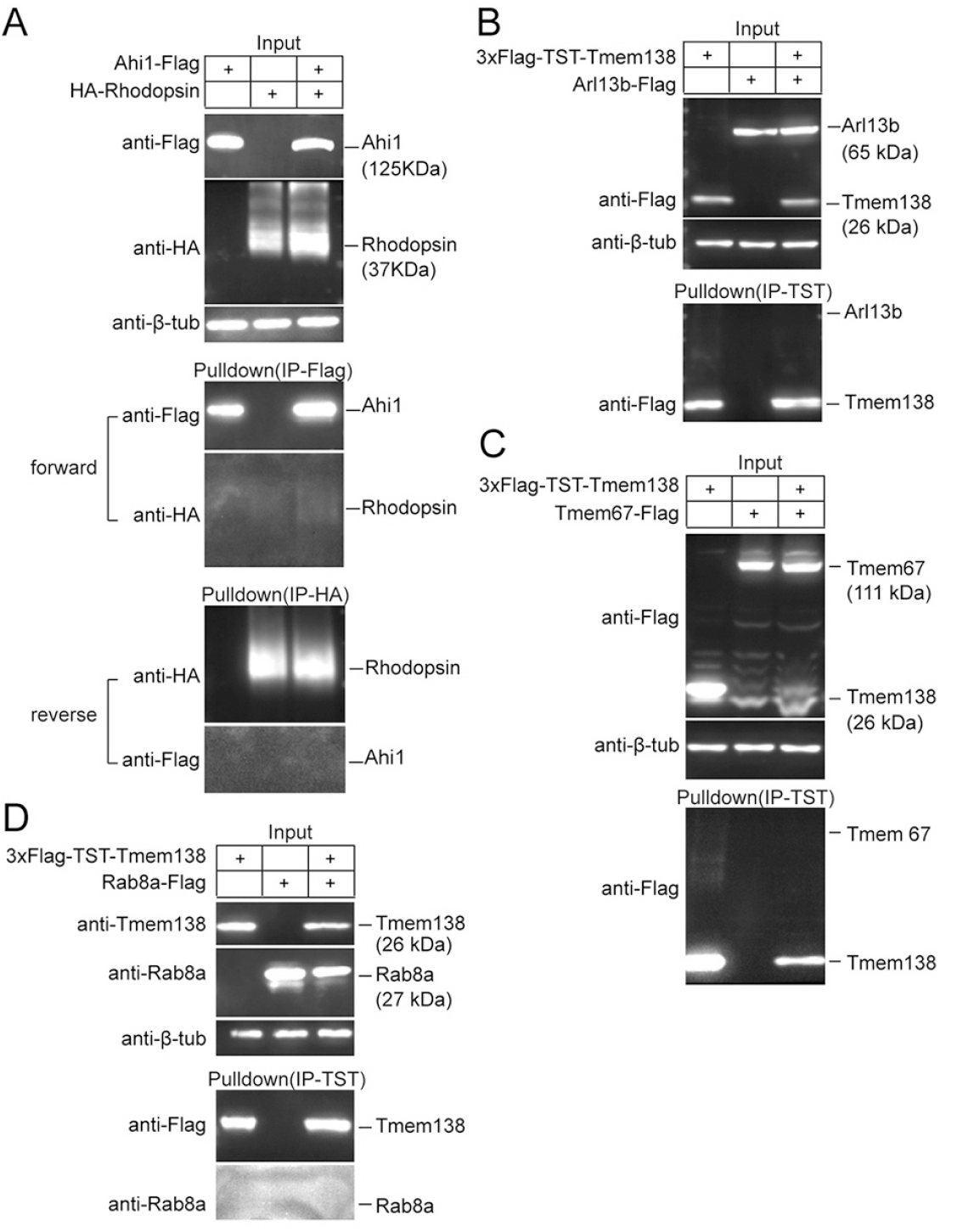
(A) Ahi1 did not interact with rhodopsin by protein pull-down assay in HEK293 cells in either forward or reverse directions. (B-D), Tmem138 did not interact with Arl13b, Tmem67, or Rab8a.

**Supplemental Figure 9.**
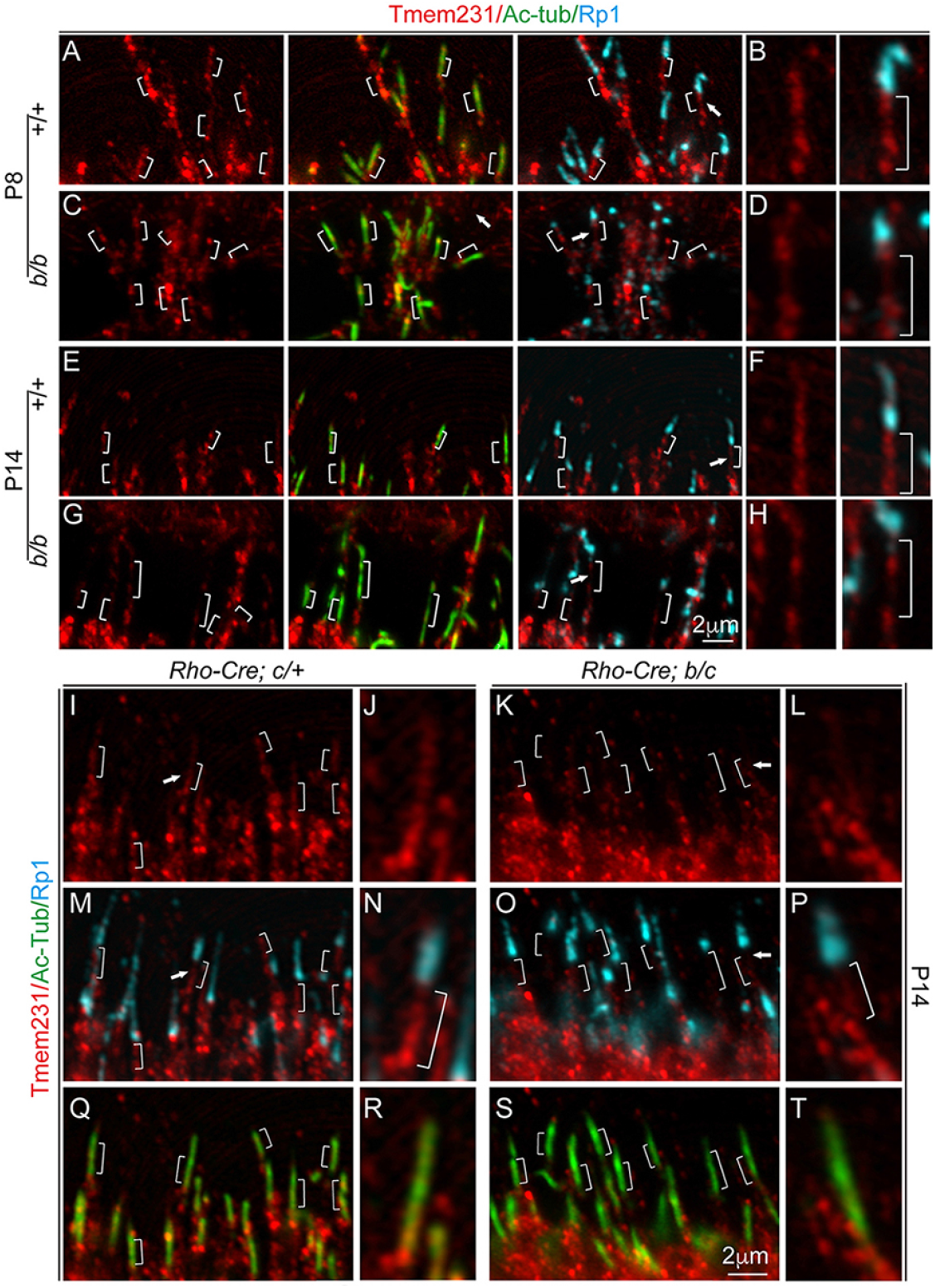
Triple labeling of Tmem231, Ac-tubulin, and Rp1. (A-H), Samples from germline knockouts and controls with indicated genotypes. (A-D) P8 retinal sections. (E-H) P14 retinal sections. (I-T), P14 retinas from conditional mutants and controls with indicated genotypes. Brackets indicate the CC domain of photoreceptors. Arrows point to the magnified cilia shown in the right.

**Supplemental Figure 10.**
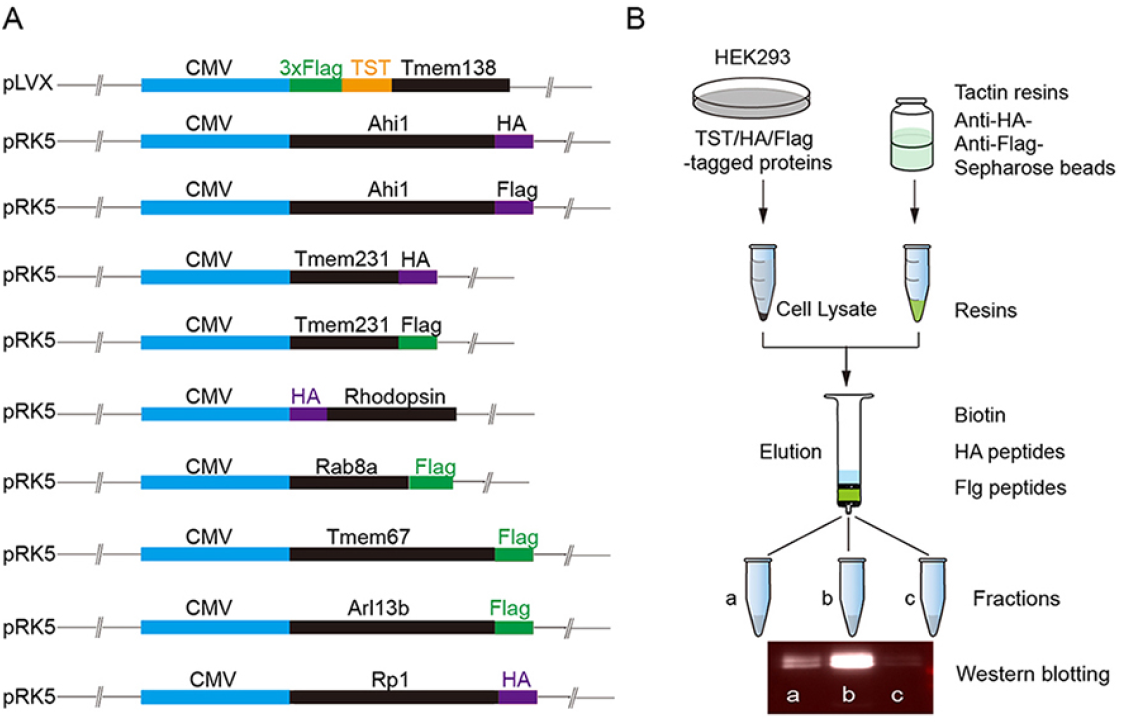
(A) DNA constructs used for detection of protein interactions in HEK293 cells. (B), Schematic illustration of major steps of affinity column pulldown experiment.

